# Coaching ribosome biogenesis from the nuclear periphery

**DOI:** 10.1101/2024.06.21.597078

**Authors:** Yinyin Zhuang, Xiangfu Guo, Olga V. Razorenova, Christopher E. Miles, Wenting Zhao, Xiaoyu Shi

**Author notes:** Correspondence (X.S.), (W.Z.).

## Abstract

Severe invagination of the nuclear envelope is a hallmark of cancers, aging, neurodegeneration, and infections. However, the outcomes of nuclear invagination remain unclear. This work identified a new function of nuclear invagination: regulating ribosome biogenesis. With expansion microscopy, we observed frequent physical contact between nuclear invaginations and nucleoli. Surprisingly, the higher the invagination curvature, the more ribosomal RNA and pre-ribosomes are made in the contacted nucleolus. By growing cells on nanopillars that generate nuclear invaginations with desired curvatures, we can increase and decrease ribosome biogenesis. Based on this causation, we repressed the ribosome levels in breast cancer and progeria cells by growing cells on low-curvature nanopillars, indicating that overactivated ribosome biogenesis can be rescued by reshaping nuclei. Mechanistically, high-curvature nuclear invaginations reduce heterochromatin and enrich nuclear pore complexes, which promote ribosome biogenesis. We anticipate that our findings will serve as a foundation for further studies on nuclear deformation.

**Highlights:** Nuclear invaginations regulate ribosome biogenesis by physically contacting nucleoli.

High-curvature nuclear tunnels increase ribosome biogenesis.

Nanopillars reduce ribosome biogenesis by transforming high-curvature nuclear invaginations to low-curvature ones.

## Introduction

Nuclear deformation, influenced by cellular and extracellular mechanical forces, plays a crucial role in human health by regulating cell fate and functions^1^. How nuclear deformation impacts cell functions is an intricate question because it appears in both healthy and diseased contexts. On the one hand, nuclear deformation is a benign morphological characteristic commonly observed in various cell types, such as stem cells and neutrophils^2–4^. On the other hand, severe nuclear deformations, such as deep nuclear invaginations, are pathological hallmarks associated with cancers^5^, neurodegeneration^6^, progeria^7^, normal aging^8^, and viral infections^9^. Therefore, dissecting the structure-function relationship in nuclear deformation is crucial to understanding when it is physiological or pathological. Closing this knowledge gap in cell biology will provide new insights in disease mechanisms and aging processes.

It is well-recognized that nuclear deformation influences mechanosensing and chromatin organization^1^. Yet, recent studies reported the elevation of ribosome biogenesis in cells with highly deformed nuclei. Buchwalter *et al*. detected abnormally high rates of ribosomal RNA (rRNA) synthesis in the nucleolus, a nuclear organelle known as a ribosome production factory, of precarious and normal aging cells^10^. These cells are notorious for their highly deformed nuclei. Similarly, increased ribosome biogenesis is a hallmark of cancer cells^11^ and embryonic stem cells ^12^, which also have severe nuclear invagination. These observations across aging, cancer, and different cell types imply a generic correlation between the invaginated nucleus and increased nucleolar activity. Since little is known about the interactions between nuclear invaginations and nucleolus, it is important to understand whether and how ribosome biogenesis can be regulated by nuclear deformation.

Electron microscopy and confocal images showed that nuclear invaginations could contact nucleoli^13–15^. Although these studies didn’t show the functional consequences of the contact, they imply potential interactions between nuclear imaginations and nucleoli. Here, we aim to identify the functional outcomes of the physical contact between the nuclear invagination and the nucleolus. First, we imaged the nuclear morphology and the nascent rRNA in the nucleolus across a series of breast epithelial cells including cancer cells. Intriguingly, the abnormally high rRNA level was found to be associated with a specific subtype of nuclear invaginations: nuclear tunnel. This observation led us to hypothesize the existence of two structural types of nuclear invaginations: a pathogenic structure that elevates nucleolar activity and a benign structure that does not excite the nucleolus.

To test this hypothesis, we dissected the structure of nuclear invaginations and examined their impact on the nucleolar ribosome biogenesis. Classifying the structural details of nuclear invaginations requires super-resolution because the nuclear invaginations can be as narrow as 100 nm^16^. We employed Label-Retention Expansion Microscopy (LR-ExM), which our lab recently developed for deep-cell imaging with up to 5 nm resolution^17, 18^. Our findings were surprising: almost all nuclear invaginations physically contact nucleoli, but only the high-curvature invaginations elevated the ribosome biogenesis in the contacted nucleoli. The regulation of nucleolar activity is so precise that the activity of the nucleolus contacted by high-curvature nuclear invagination can be higher than that contacted by low-curvature nuclear invaginations within the same nucleus.

To quantitatively understand the structure-function relationship between the curvature of the nuclear invagination and nucleolar ribosome biogenesis, we employed a cutting-edge nanomaterial: nanopillars^19–21^. Nanopillars have been used to guide both plasma membrane curvature^22–25^ and subnuclear deformation in live cells^19–21^, as well as mimic the fibrous extracellular matrix to understand nucleus deformation during cell migration^26^. We induced nuclear invaginations with controlled curvatures in live cells, by growing cells on a substrate covered by nanopillars with specific radii. Shockingly, we found a quasi-linear relationship between the rRNA level and the nuclear invagination curvature. Based on this finding, we successfully reduced the rRNA levels of breast cancer and progeria cells by deforming their nuclei to low-curvature invaginations using wide nanopillars. The results from the nanopillar experiment are groundbreaking. These demonstrate that structural alterations to the nucleus, without direct genetic interference, can rescue nucleolar function and ribosome biogenesis. This finding not only validates our hypothesis on the structure-function relationship in nuclear invagination but also shines light on a potential therapeutic approach, distinct from traditional gene therapy or pharmaceutics.

Furthermore, we investigated the mechanisms of how nuclear invaginations regulate nucleolar activity and what causes nuclear invaginations to form. These mechanisms involve complex interactions among organelles and structures, including nuclear lamina, chromatin, nuclear pore complexes (NPCs), ribosome exporters, cytoskeleton, and possibly endoplasmic reticulum (ER) and mitochondria. In the scope of this study, we focus on the interactions inside the nucleus and near the nuclear envelope (NE). We anticipate our findings to be a starting point for more sophisticated studies of the functions of deformed NE achieved through other organelle-organelle interactions.

In the upcoming section, we provide experimental evidence on the structure-function relationship between nuclear invagination and ribosome biogenesis. Our exploration began with the association of nuclear invaginations with nucleolar activity. We observed frequent physical contact between the NE and the nucleolus in cultured cells and human tissues. Notably, the nuclear tunnel—a high-curvature type of invagination—emerges as a key enhancer of rRNA synthesis and pre-ribosome assembly in the nucleolus. This curvature-dependent regulation offers a gateway to manipulating ribosome biogenesis, as evidenced by using nanopillars to lower invagination curvature, which effectively reduces ribosome production in progeria cells and breast cancer cells. Further, the narrative will unpack the mechanism of how nuclear invaginations interact with other organelles to orchestrate ribosome biogenesis. High-curvature nuclear invaginations attenuate heterochromatin, enrich NPCs, and spatially arrange other organelles in its proximity, which all favor ribosome biogenesis. To elucidate the biophysical picture behind the regulatory mechanism, we designed a diffusion model that explains how heterochromatin enrichment inversely affects ribosome biogenesis. The final part of the results provides a preliminary understanding of the causes of nuclear deformation, opening more possibilities to regulate ribosome biogenesis.

## Results

### Structure-function relationship between nuclear invagination and the nucleolus

#### Nuclear invagination is associated with ribosome biogenesis

The first question we addressed was whether there is a correlation between the nuclear invagination level and the ribosome biogenesis rate at the single-cell level. We conducted a study where we fluorescently co-labeled the NE and nascent rRNA in a series of breast epithelial cell lines ranging from immortalized (MCF-10A) to cancerous exhibiting a gradient of nuclear invagination from subtle to severe^21, 27, 28^. The NE was outlined using immunostaining of lamin B2 (Figure 1A, i-v), while fluorescence from 5-Ethynyl Uridine (EU) highlighted pulse-labeled nascent RNAs (Figure 1A, vi-x). Since rRNAs make up the majority of cellular RNAs, we used the EU signal within the nucleolus (Figure 1Avi, white arrowheads) as a proxy for nucleolar activity in ribosome biogenesis^10^. By comparing the level of nuclear invagination with EU intensity, we aimed to determine if there is a correlation between nuclear invagination and ribosome biogenesis.

**Figure 1.**
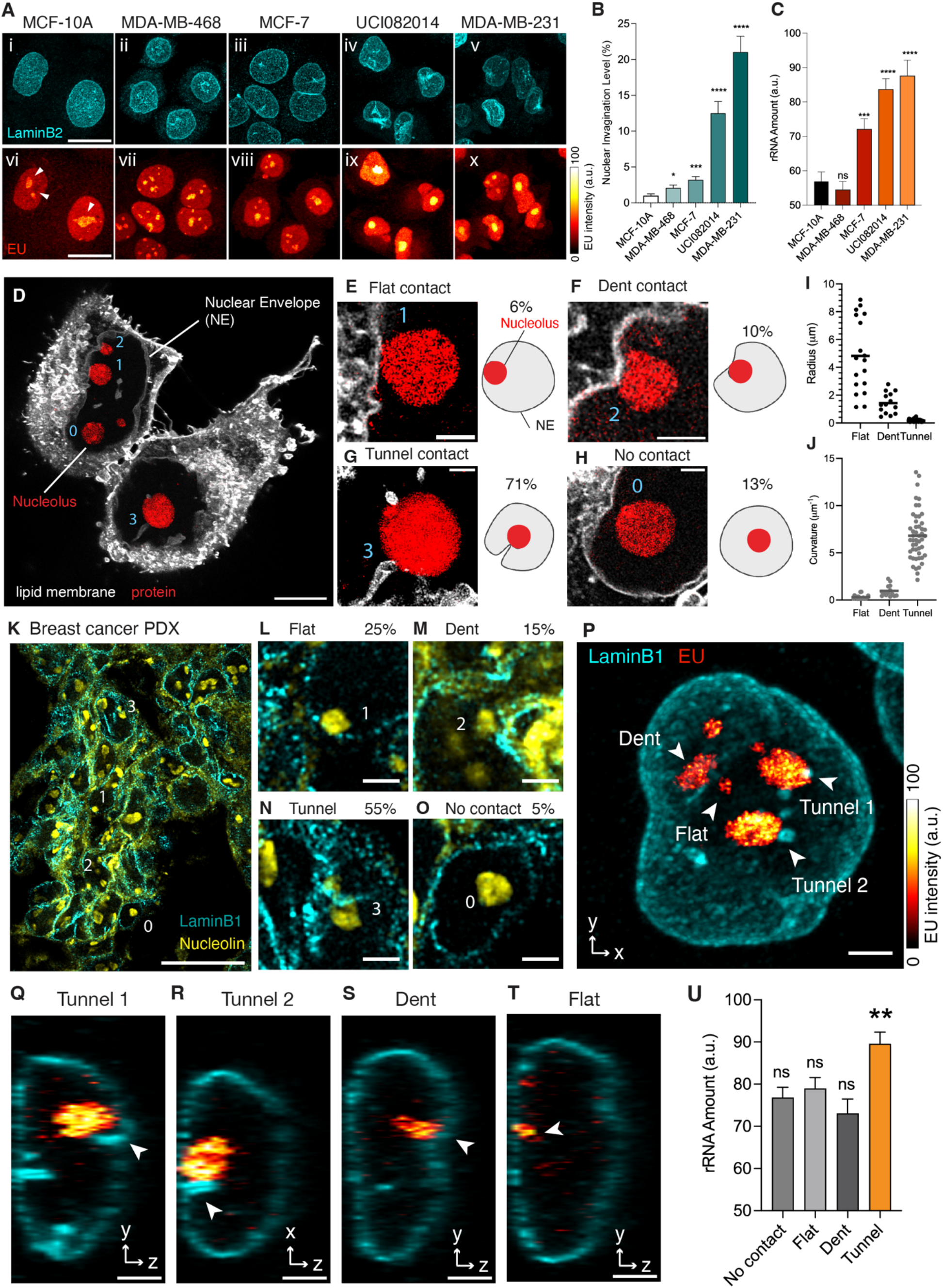
Nuclear Invagination-Nucleolus Contact Regulates Ribosome Biogenesis. (A) Images of breast cell lines pulse-labeled with EU (red hot) for 1 hour followed by immunostaining with anti-LaminB2 antibody (cyan). Images i-v and vi to x: MCF-10A, MDA-MB-468, MCF-7, UCI082014, and MDA-MB-231 cells. Arrowheads point to nucleoli. Color bar: EU intensity from 0 to 100. Scale bar: 20 µm. (B) Barchart of nuclear invagination levels of MCF-10A, MDA-MB-468, MCF-7, UCI082014, and MDA-MB-231 cells. The invagination level is measured as the percentage of folded area out of the total area of the nuclear envelope, in the maximum intensity projection of three-dimensional images. The nuclear envelope is maintained with anti-LaminB2 antibody. Each bar represents the mean ± standard error of more than 30 nuclei of each cell line from 3 independent experiments. * indicates p<0.05, *** indicates p<0.001, and **** indicates p<0.0001 compared to MCF-10A cells by Welch’s t test. (C) Barchart of newly synthesized rRNA in the nucleoli of MCF-10A, MDA-MB-468, MCF-7, UCI082014, and MDA-MB-231 cells. The rRNA amount is measured as the EU intensity within each nucleolus in the maximum intensity projection of three-dimensional EU Airyscan images. Each bar represents the mean ± standard error of more than 30 nucleoli of each cell line from 3 independent experiments. *** indicates p<0.001, **** indicates p<0.0001 and ns indicates p>0.05 compared to MCF-10A cells by Welch’s t test. (D) Expansion microscopy image of UCI082014 cells stained with lipid (white) and protein (red) dyes. 0, 1, 2, and 3 indicate no, flat, dent, and tunnel-type NE-nucleolus contact, respectively. Length expansion factor: 3.9. Scale bar: 5 µm in pre-expansion unit. (E-H) Left: Zoom-in images of different NE-nucleolus contacts from (D). Scale bar: 1 µm in pre-expansion unit. Right: Illustration of different types of NE-nucleolus contact. The percentage of nucleoli contacted by flat NE, dents, tunnels, and without contact is 6%, 10%, 71%, and 13%, respectively. Total number of nucleoli analyzed: 103. (I) Distribution of radii of flat NE, nuclear dents, and nuclear tunnels. The scatter plot represents the radius of individual structures with mean ± standard error from 18 flat areas, 14 dents, and 49 tunnels. (J) Scatter plot of the curvature of flat NE, nuclear dents, and nuclear tunnels converted from (I). (K) Images of tumors from breast cancer patient-derived xenograft immunostained with anti-LaminB1 (cyan) and anti-nucleolin (yellow) antibodies. 0, 1, 2, and 3 indicate no, flat, dent, and tunnel type NE-nucleolus contact, respectively. Scale bar: 20 µm. (L-O) Zoom-in images of different types of NE-nucleolus contacts from (K). Scale bar: 2 µm. The percentage of nucleoli contacted by NE flat, dents, tunnels, and without contact is 25%, 15%, 55%, and 5%, respectively. Total number of nucleoli analyzed: 67. (P) Image of an MCF-7 cell pulse-labeled with EU (red hot) for 1 hour, then immunostained with anti-LaminB1 antibody (cyan). Arrowheads point to nucleoli contacted by flat NE, nuclear dents, or nuclear tunnels. Color bar: EU intensity from 0 to 100. Scale bar: 3 µm. (Q-T) Side views of (P). Arrowheads point to nucleoli contacted by flat NE, nuclear dents or nuclear tunnels. Scale bar: 3 µm. (U) Barchart of newly synthesized rRNA in individual nucleolus without NE contact or contacted by flat NE, nuclear dents, or nuclear tunnels. Each bar represents mean ± standard error of more than 30 nucleoli. ** indicates p<0.01 and ns indicates p>0.05 by unpaired t test. All images in this figure were taken by an Airyscan microscope with a 63x objective.

In Figure 1A, breast cell lines are ordered from left to right based on the severity of nuclear invagination, ranging from subtle to severe (Figures 1Ai-v, and 1B). Correspondingly, the amount of rRNA in the nucleoli increased from left to right panels (Figures 1Avi-x, and 1C). This trend indicates a positive correlation between nuclear invagination levels and rRNA amounts. Since rRNA is only one component of the ribosome, we also examined other ribosomal components recruited after rRNA synthesis. These include the pre-60S ribosomal subunit eIF6, recruited during pre-ribosome assembly in the nucleolus^29^, and the ribosomal protein RPL13, recruited during ribosome maturation in the cytoplasm^30^. The imaging and western blotting results showed that the amounts of both eIF6 and RPL13 are positively correlated with nuclear invagination levels in the breast cancer cell lines (Figure S1). The consistent trends in rRNA, pre-ribosome, and ribosome production suggest that the ribosome biogenesis rate positively correlates with the level of nuclear invagination.

#### Three types of physical contact between NE and nucleoli

Next, we examined the structure and dynamics of nuclear invaginations and their spatial relationship with nucleoli. Studying nuclear invaginations is challenging due to their narrow diameter, sometimes as small as 100 nm^16^, necessitating super-resolution microscopy. Additionally, multiplex imaging is required to visualize the spatial relationship between nuclear invaginations and nucleoli. To meet these needs, we developed an expansion microscopy method combining LR-ExM from our lab^17, 18^ and expansion microscopy protocols from other labs^31–33^. This approach enabled us to analyze hundreds of cells in 3D (Movie S1), leading to a series of surprising findings in the spatial relationships between the NE and nucleoli with statistical rigor.

Our first finding revealed a high probability of NE-nucleolus contact in cells and tissues. Contrary to textbook depictions of nucleoli as suspended spheres, our 3D images of triple-negative breast cancer cells (UCI082014) showed that approximately 87% of nucleoli were in contact with or adjacent to the NE, with only about 13% having no NE contact (Figures 1D-H, Movie S2). This high contact probability was also observed in tumors formed from breast cancer patient-derived xenograft (PDX-HCI-002) (Figures 1K-O) and various cell types, including immortalized breast epithelial cells (MCF-10A), primary cells from progeria patients, mouse embryonic fibroblasts (MEF), and human osteosarcoma cells (U2OS). The contact probability between NE and nucleoli varied among cell lines but was always over 80% (Figure S2).

We classified NE-nucleolus contact into three types based on the NE curvature at the contact site: flat, dent, and tunnel contacts. The flat contact is the contact between smooth NE and a nucleolus (Figure 1E). The radius of a flat contact site ranges from 1 µm to 9 µm (Figure 1I), with an average curvature of 0.31 µm^-1^, which is close to the overall curvature of a nucleus (Figure 1J). The dent contact (Figure 1F) represents the contact between a dent-shaped nuclear invagination and a nucleolus. A nuclear dent has a radius larger than 200 nm (Figure 1I) and a curvature lower than 1 µm^-1^ (Figure 1J). It is also termed as a nuclear indentation in literature^34, 35^. Our measurements show that the radius of a nuclear dent ranges from 0.5-3 µm (Figure 1I), with an average curvature of 0.96 µm^-1^ (Figure 1J). The tunnel contact occurs between a tunnel-shaped nuclear invagination and a nucleolus (Figure 1G). The nuclear tunnels have radii ranging from 50-200 nm (Figure 1I). Their average curvature is 6.82 µm^-1^, significantly higher than the curvature of dents or smooth NE (Figure 1J, Movie S1). Nuclear tunnels are also termed nucleoplasmic reticulum or deep nuclear invagination in literature^16, 36^. These three types of NE-nucleolus contact were observed not only in cultured cells but also in PDX-based tumors, indicating their pathological relevance (Figures 1K-N).

#### Dynamics of the nuclear invagination and its contact with the nucleolus

To evaluate the stability of NE-nucleolus contact, we tracked the three types of NE-nucleolus contacts over cell cycles. We used a CRISPR knock-in cell line expressing mNeongreen-upstream binding transcription factor (UBTF) at its endogenous level, where the mNeongreen signal marks the position of the nucleolus^37^. ERtracker live cell staining was used to indicate the nuclear invaginations, in the same way as demonstrated in previous studies^38^. The live cell video showed that the tunnel-nucleolus contact can last through the whole interphase until mitosis (Movies S3&S4 and Figure S3A). In contrast, flat and dent contacts are more transient than tunnel contact. The dent-nucleolus contact lasted for about half an hour (Movie S5 & Figure S3B). Given the differences in dynamics and structure of these NE-nucleolus contacts, we questioned whether the various types play different roles in regulating ribosome biogenesis.

#### Only the nuclear tunnel increases nucleolar activity

The study’s most significant finding is that nucleolar activity depends on the type of nuclear invagination it contacts. This dependency is so localized that, within a single nucleus, individual nucleoli contacted by different types of nuclear invaginations exhibit distinct activities (Figure 1P). Nucleoli in contact with nuclear tunnels showed abundant rRNAs (Figures 1P, Q, and R), while those in contact with nuclear dents or flat NE showed fewer rRNAs (Figures 1P, S, and T). This precise regulation suggests that the controlling factor is a local structure or distribution. Statistically, contact with nuclear tunnels is associated with increased nucleolar activity in rRNA synthesis by about 20% compared to that without contact, whereas contact with dents or flat NE does not alter nucleolar activity (Figure 1U).

Despite the correlation between nuclear tunnels and high nucleolar activity, it is unclear whether the nuclear tunnel is the cause or result of an active nucleolus. To address this question, we needed a tool to initiate nuclear invaginations directly, ideally without gene editing or drug treatments that could affect the nucleolus. The nanopillar substrate for cell culture offers an excellent solution^19, 21^. This glass surface is fabricated with arrays of vertical pillar-like structures with designed radii, height, and pitch (Figures 2A and B). When cells grow on these substrates (Figure 2C), the nanopillars deform the nuclei and generate artificial nuclear invaginations in live cells (Figure 2D). These artificial invaginations closely resemble natural nuclear tunnels (Figure 2E). If these artificial tunnels increase rRNA synthesis in nucleoli, it would suggest that nuclear tunnels cause elevation of nucleolar activity, which is exactly what our nanopillar experiments demonstrated.

**Figure 2.**
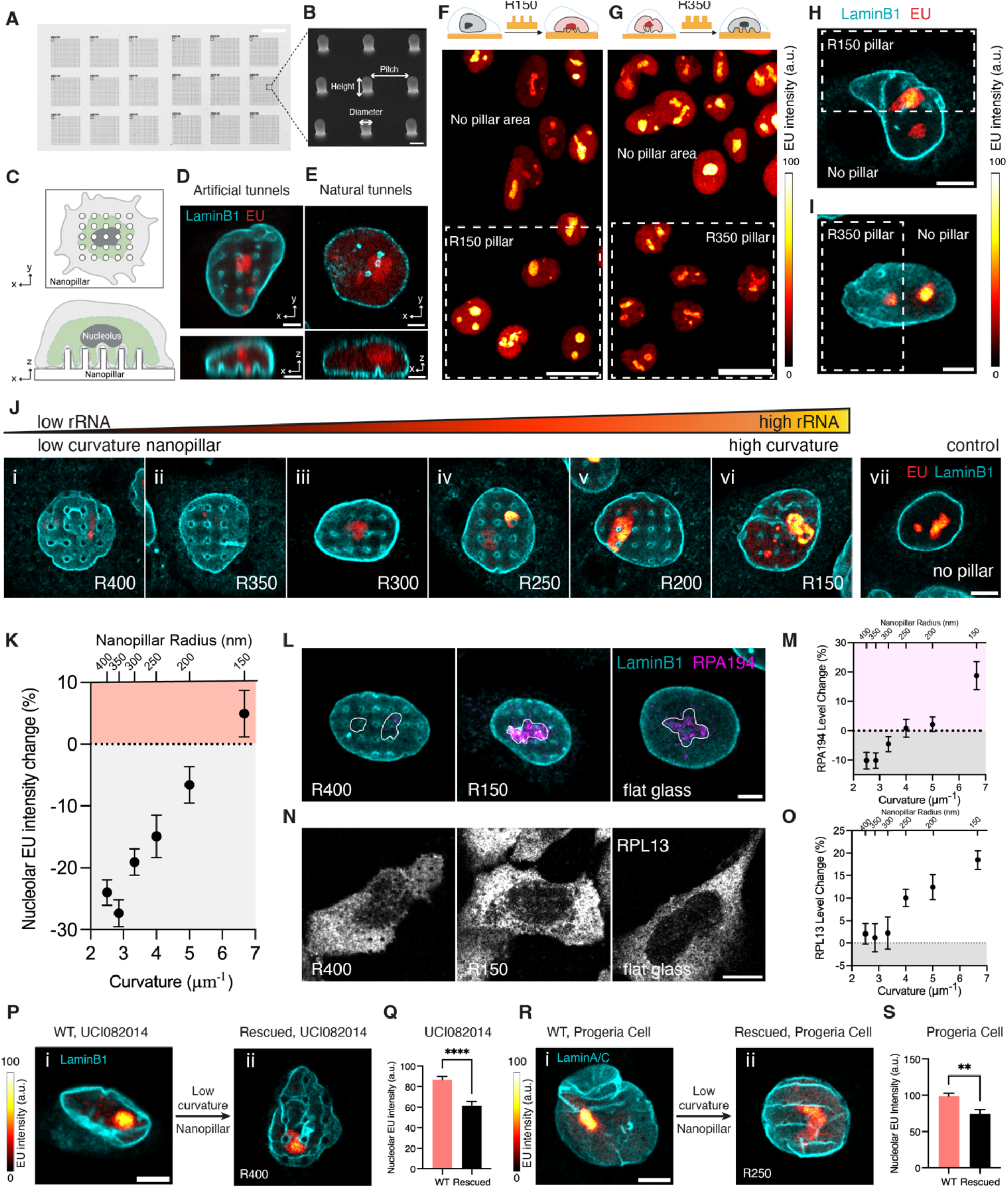
Regulation of Nucleolar Activity is Curvature-Dependent. (A) Brightfield image of nanopillar patterns with different radii. Scale bar: 20 µm. (B) Scanning electron microscopy image of 9 nanopillars with 3 µm pitch and 1.5 µm height. Scale bar: 1 µm. (C) Illustration of top and side views of a cell cultured on a nanopillar substrate. (D) Image of top and side views of MCF-7 cells cultured on nanopillars with a radius of 200 nm, pulse-labeled with EU (red) for 1 hour, and immunostained with anti-LaminB1 antibody (cyan). Scale bar: 3 µm. (E) Image of top and side views of MDA-MB-468 cell with natural nuclear tunnels. The cell was pulse-labeled with EU (red) for 1 hour and immunostained with anti-LaminB1 antibody (cyan). Scale bar: 3 µm. (F) Upper schematics: Illustration of an MCF-10A cell seeded on nanopillars with a radius of 150 nm. Bottom: Image of MCF-10A cells seeded on nanopillars with a radius of 150 nm (dashed line box) and compared to the cells on the flat region without nanopillars. The cells were pulse-labeled with EU (red hot) for 1 hour. Color bar: EU intensity from 0 to 100. Scale bar: 20 µm. (G) Upper schematics: Illustration of an MCF-10A cell seeded on nanopillars with a radius of 350 nm. Bottom: Image of MCF-10A cells seeded on nanopillars with a radius of 350 nm (dashed line box) and compared to the cells on the flat region without nanopillars. The cells were pulse-labeled with EU (red hot) for 1 hour. Color bar: EU intensity from 0 to 100. Scale bar: 20 µm. (H) Image of an MCF-10A that has half the nucleus on the nanopillars with a radius of 150 nm (dashed line box) and half the nucleus on flat glass. The cell was pulse-labeled with EU (red hot) for 1 hour. Color bar: EU intensity from 0 to 100. Scale bar: 5 µm. (I) Image of an MCF-10A that has half the nucleus on the nanopillars with a radius of 350 nm (dashed line box) and half the nucleus on flat glass. The cell was pulse-labeled with EU (red hot) for 1 hour. Color bar: EU intensity from 0 to 100. Scale bar: 5 µm. (J) Images of MCF-10A cells seeded on nanopillars with gradient radii from 400 nm to 150 nm (i to vi) and on flat glass without nanopillars (vii). The cells were pulse-labeled with EU (red hot) for 1 hour and then immunostained with anti-LaminB1 antibody (cyan). Color bar: EU intensity from low to high. Scale bar: 5 µm. (K) Scatter plot of the amount change of newly synthesized rRNA in the nucleoli of MCF-10A cells on nanopillars with different radii from 400 nm to 150 nm, normalized to that of cells on flat no pillar region. Each data point represents the mean ± standard error of more than 100 cells from 3 independent experiments. (L) Images of MCF-10A cells seeded on nanopillars with radius of 400 nm or 150 nm, or on flat no pillar region. The cells were immunostained with anti-LaminB1 (cyan) and anti-RPA194 (magenta) antibodies. The white lines outline the nucleoli. Scale bar: 5 µm. (M) Scatter plot of the amount change of RPA194 in the nucleoli of MCF-10A cells on nanopillars with different radii from 400 nm to 150 nm, normalized to that of cells on flat glass. Each data point represents the mean ± standard error of more than 60 cells from 3 independent experiments. (N) Images of MCF-10A cells seeded on nanopillars with a radius of 400 nm or 150 nm or on flat glass. The cells were immunostained with anti-RPL13 antibodies (grey). Scale bar: 10 µm. (O) Scatter plot of the amount change of RPL13 in MCF-10A cells on nanopillars with different radii from 400 nm to 150 nm, normalized to that of cells on flat glass. Each data point represents the mean ± standard error of more than 30 cells from 3 independent experiments. (P) i: Image of a UCI082014 cell cultured on flat surface. ii: Image of a UCI082014 cell cultured on nanopillars with 400 nm radius. Both cells were pulse-labeled with EU (red hot) for 1 hour then immunostained with anti-LaminB1 antibody (cyan). Color bar: EU intensity from 0 to 100. Scale bar: 5 µm. (Q) Barchart of the amount of newly synthesized rRNA in the nucleoli of UCI082014 cells cultured on a flat surface and on nanopillars with 400 nm radius. Each bar represents the mean ± standard error of more than 40 cells from 3 independent experiments. **** indicates p<0.0001 by unpaired t test. (R) i: Image of an HGPS cell cultured on a flat surface. ii: Image of an HGPS cell cultured on nanopillars with 400 nm radius. Both cells were pulse-labeled with EU (red hot) for 1 hour then immunostained with anti-LaminB1 antibody (cyan). Color bar: EU intensity from 0 to 100. Scale bar: 5 µm. (S) Barchart of the amount of newly synthesized rRNA in the nucleoli of HGPS cells cultured on a flat surface and on nanopillars with 400 nm radius. Each bar represents the mean ± standard error of more than 40 cells from 3 independent experiments. **** indicates p<0.0001 by unpaired t test. All fluorescence images in this figure were taken by an Airyscan microscope with a 63x objective.

We first fabricated nanopillar arrays with a radius of 150 nm (R150 nanopillar), matching the average radius of natural nuclear tunnels (Figure 1I). We seeded MCF-10A cells on the R150 nanopillar substrate (Figure S4A). MCF-10A cells, which are immortalized breast epithelial cells, have the fewest nuclear invaginations (Figure 1Ai) and the lowest ribosome biogenesis level (Figure 1Avi). Once the cells adhered to the nanopillars, we stained newly synthesized RNAs with EU and later labeled the NE with an anti-Lamin B1 antibody. The super-resolution images of these nanopillar-treated cells were very exciting. The rRNA levels in nucleoli contacted by nanopillar-induced nuclear tunnels were higher than in those on the flat area of the same substrate (Figure 2F). Like the natural case (Figure 1P), the impact of a nanopillar-induced nuclear tunnel is spatially confined to the nucleolus that the tunnel contacts. Figure 2H illustrates a cell with half of the nucleolus on the R150 nanopillar and the other half on flat glass, providing further evidence of local activation of the nucleolus at the sub-nucleus level. The nucleolus contacted by R150-induced nuclear tunnels contained more rRNA than the nucleolus in the other half nucleus on flat glass (Figure 2H). These results confirmed that nuclear invaginations can cause changes in ribosome biogenesis.

Curiously, we also fabricated low-curvature nanopillar arrays with a 350 nm radius (R350 nanopillar), matching the radius and curvature of natural nuclear dents which should not promote nucleolus activity. Imaging MCF-10A cells cultured on R350 nanopillars, we found that nucleoli contacted by R350-induced nuclear dents produced less rRNA than those on the flat area of the same substrate (Figure 2G). In a cell with half of the nucleolus on the R350 nanopillar and the other half on flat glass, the nucleolus contacted by R350-induced nuclear dents contained fewer rRNAs than the nucleolus in the other half nucleus on flat glass (Figure 2I). These results suggest that the relatively low nucleolar activity of non-cancer cells can be further reduced by nanopillar-induced nuclear dents. We suspect this suppression occurs because the R350 nanopillars converted natural tunnels to nuclear dents (Figure S4B). The R150 and R350 experiments show that nanopillars can both upregulate and downregulate nucleolar activity by fine-tuning the curvature of nuclear invaginations.

#### Curvature-dependent regulation of nucleolar activity

The nanopillar substrate enables a quantitative investigation of how nuclear invaginations with different curvatures influence nucleolar activity. We engineered a substrate with nanopillar arrays that precisely generate nuclear invaginations with a range of curvatures. These nanopillars are uniform in radius within each array but vary between arrays across the substrate from 150 nm to 400 nm in a graded manner (Figure S5). We observed that rRNA synthesis significantly increased as the curvature of nanopillar-induced nuclear invaginations ascended (Figure 2J). Specifically, rRNA synthesis was upregulated in MCF-10A cells on high-curvature nanopillars with a radius of 150 nm (Figure 2Jvi) compared to control cells growing on flat glass (Figure 2Jvii). Conversely, rRNA synthesis was downregulated in cells on low-curvature nanopillars with radii ranging from 250 to 400 nm (Figure 2Ji-iv). The threshold radius between upregulation and downregulation was around 200 nm, corresponding to a curvature of 5 µm⁻¹. Intriguingly, Figure 2K shows a linear positive correlation between nucleolar activity and nanopillar curvature. In this plot, R350-induced nuclear invaginations decreased rRNA levels by an average of 27% and up to 73%, while R150-induced nuclear invaginations increased rRNA levels by an average of 5% and up to 80%. The efficiency of upregulation was lower than that of downregulation because MCF-10A cells had intrinsic nuclear tunnels that elevated the baseline of rRNA synthesis (Figure S4B, arrows). The overall 30% change in rRNA synthesis caused by nanopillars is significant, matching the difference between cells derived from normal breast epithelium and aggressive triple-negative breast cancers (Figure 1C). Our results demonstrate that the regulation of nucleolar activity is dependent on the nuclear invagination curvature.

In addition to rRNA, we examined the impact of invagination curvature on other markers of ribosome biogenesis, including RPA194, an RNA polymerase I subunit responsible for rDNA transcription, and RPL13, a key component of mature ribosomes. We found an increase in RPA194 (Figures 2L and M) and RPL13 (Figures 2N and O) in cells cultured on high-curvature nanopillars. We concluded that the curvature of nuclear invagination is crucial for regulating ribosome biogenesis. The higher the invagination curvature, the greater the amount of rRNA and pre-ribosomes produced by the contacted nucleolus.

#### Low-curvature nanopillars reduce ribosome biogenesis in progeria and cancer cells

Now, we established a curvature-dependent relationship between nuclear invagination and nucleolar activity. Based on this structure-function relationship, we hypothesize that reshaping the nuclei of diseased cells can rescue the overactive ribosome biogenesis observed in these cells. To test this hypothesis, cancer and progeria cells were cultured on substrates with low-curvature nanopillars. According to the data presented (Figures 2K, M&O), nanopillars with a radius larger than 250 nm are classified as low-curvature pillars and are effective in decreasing ribosome biogenesis.

We applied these low-curvature nanopillars to a cell line derived from a primary tumor of a triple-negative breast cancer patient (UCI082014) and primary fibroblasts from a patient with Hutchinson-Gilford progeria syndrome (HGPS). Although these cell types harbor different mutations, they share similar pathological phenotypes, including abundant high-curvature nuclear invaginations and elevated ribosome biogenesis^10, 28^. These intrinsic high-curvature nuclear invaginations manifest as single tunnels and sharp folds, which are considered connected tunnels (Figures 2Pi&Ri). During cell culturing, the intrinsic high-curvature invaginations were transformed into nuclear dents by the low-curvature nanopillars. Remarkably, this morphological change led to a reduction in rRNA levels in the diseased cells to a physiological level (Figures 2P-S). The low-curvature nanopillars effectively reduced ribosome biogenesis in both breast cancer and progeria cells, despite the distinct etiologies of these diseases. This result suggests that precise control of nanoscale nuclear membrane curvature can be a general strategy for regulating ribosome biogenesis.

### Cellular mechanisms of curvature-dependent regulation of ribosome biogenesis

In the previous section, we showed that the high-curvature nuclear invaginations activate ribosome biogenesis, while low-curvature nuclear invaginations have negligible impact. Below we explored mechanisms of this curvature-dependent regulation of ribosome biogenesis. Our major approach was the 3D multicolor expansion microscopy. We imaged the lipid membrane, nucleoli, chromatin, and other organelles and proteins with our expansion microscopy method, which clearly displayed the detailed spatial relationships between the nuclear invaginations and other components known to contribute to ribosome biogenesis.

#### Nuclear tunnels interact with multiple organelles to facilitate ribosome biogenesis

Our first finding in the mechanism is that the high-curvature nuclear invaginations interact with chromatin and organelles involved in ribosome biogenesis. Ribosome biogenesis involves multiple organelles and hundreds of proteins^39–41^. It begins in the nucleolus, where chromatin unfolds to expose ribosomal DNA (rDNA) for transcription within the fibrillar center (FC)/dense fibrillar component (DFC) of the nucleolus. The presence of FC/DFC regions is a structural hallmark of active ribosome biogenesis. In these regions, rRNAs are synthesized by RNA polymerase I (polI). Following the rRNA synthesis, rRNA processing occurs in the DFC, and the initial steps of pre-ribosome subunit assembly in the granular component (GC)^37, 39^. These pre-ribosome subunits then bind with exportin1 in the nucleoplasm^42, 43^. Exportin1-bound pre-ribosomes pass through the nuclear pore complexes (NPCs) to the cytoplasm, where they undergo final assembly and modification to become fully functional ribosomes^29, 44^. Although ER is not involved in their biogenesis, ribosomes associate with the ER to synthesize proteins. We found that high-curvature nuclear invaginations spatially organize all these organelles on both sides of the NE.

On the nuclear side, we observed that the nucleoli developed more active FC and DFC regions when contacted by nuclear tunnels. Figure 3A shows that the nucleoli in contact with tunnels have developed FC/DFC regions. We immunostained RPA194, the largest subunit of polI. The colocalization between RPA194 and FC/DFC regions confirmed active rRNA synthesis in the nucleoli contacted by nuclear tunnels (Figure 3B). In contrast, uncontacted nucleoli or those contacted by nuclear dents had much fewer RPA194 and fewer FC/DFC regions, indicating lower rRNA synthesis rates (Figure 3C). Statistically, nucleoli contacted by tunnels had 83% more active FC/DFC regions than those uncontacted (Figure 3D).

**Figure 3.**
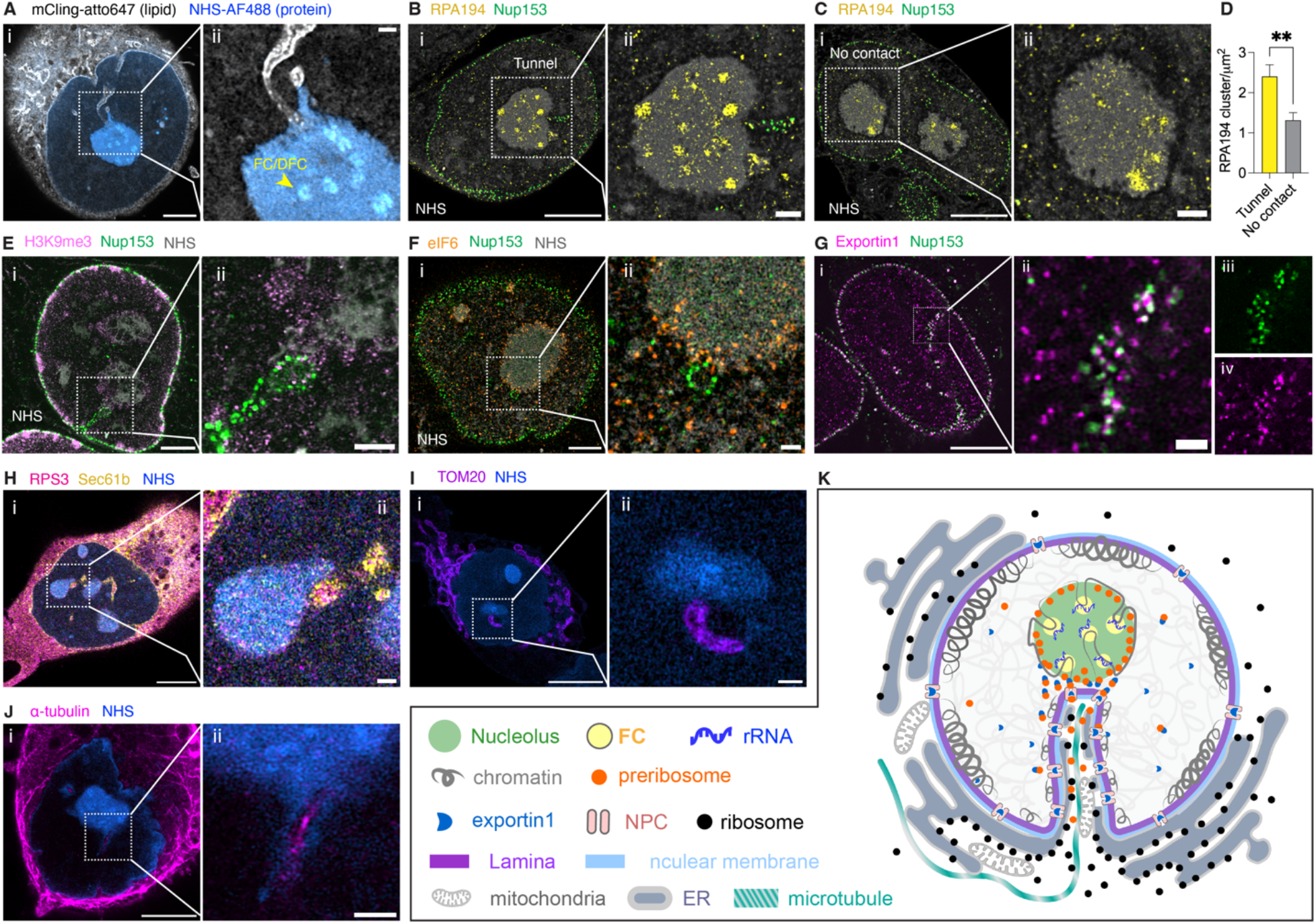
Mechanisms of Nuclear Tunnels in Organizing Cellular Structures for Ribosome Biogenesis. All fluorescence images are Expansion Microscopy images taken on an Airyscan microscope with a 63x objective. All scale bars are in pre-expansion units. (A) i: Image of total lipid and total protein of a UCI082014 cell. Lipids were stained with mCling-atto647 (grey) and proteins were stained with Alexa Fluor 488 NHS ester (blue). Scale bar: 3 µm. ii: Magnified view of nuclear tunnel-nucleolus contact in the boxed area of the image (i). The arrow points to an FC/DFC region of the nucleolus. Scale bar: 500 nm. Length expansion factor: 3.9. (B) i: Image of an active nucleolus contacted by a nuclear tunnel in a UCI082014 cell. The cell was stained with anti-RPA194 antibodies (yellow), which mark Pol I, anti-Nup153 antibodies (green), which mark the nuclear tunnel, and NHS ester (grey). Scale bar: 5 µm. ii: Magnified view of NE-nucleolus contact in the boxed area of the image (i). Scale bar: 1 µm. Length expansion factor: 3.9. (C) i: Image of an inactive nucleolus without contact with NE in a UCI082014 cell. The cell was stained with anti-RPA194 antibodies (yellow), anti-Nup153 antibodies (green), and NHS ester (grey). Scale bar: 5 µm. ii: Magnified view of NE-nucleolus contact in the boxed area of the image (i). Scale bar: 1 µm. Length expansion factor: 3.9. (D) Barchart of RPA194 cluster density in the nucleoli contacted by tunnels or suspended in UCI082014 cells. Each bar represents the mean ± standard error of more than 15 nucleoli from 3 independent experiments. ** indicates p<0.01 by unpaired t test. (E) i: Image of an MCF-10A cell stained with anti-H3K9me3 antibodies (pink), anti-Nup153 antibodies (green), and NHS ester (grey). Scale bar: 3 µm. ii: Magnified view of tunnel-nucleolus contact in boxed area of image (i). Scale bar: 1 µm. Length expansion factor: 3.9. (F) i: Image of a UCI082014 cell stained with anti-eIF6 (orange) antibodies, anti-Nup153 (green), and NHS ester (grey). Scale bar: 3 µm. ii: Magnified view of tunnel-nucleolus contact in boxed area of image (i). Scale bar: 500 nm. Length expansion factor: 3.7. (G) i: Image of a UCI082014 cell stained with anti-Nup153 antibodies (green) and anti-exportin1 (magenta) antibodies. Scale bar: 5 µm. ii: Magnified view of a nuclear tunnel in boxed area of image (i). Scale bar: 500 nm. iii: Magnified view of Nup153 channel of image (ii). iv: Magnified view of exportin1 channel of image (ii). Length expansion factor: 4.0. (H) i: Image of a UCI082014 cell stained with anti-Sec61b antibodies (yellow) and anti-RPS3 (pink) antibodies, and NHS ester (blue). Scale bar: 5 µm. ii: Magnified view of tunnel-nucleolus contact in boxed area of image (i). Scale bar: 500 nm. Length expansion factor: 3.9. (I) i: Image of a UCI082014 cell stained with anti-Tom20 antibodies (purple) and NHS ester (blue) and. Scale bar: 5 µm. ii: Magnified view of tunnel-nucleolus contact in boxed area of image (i). Scale bar: 500 nm. Length expansion factor: 3.7. (J) i: Image of a UCI082014 cell stained with anti-alpha-tubulin antibodies (magenta) and NHS ester (blue). Scale bar: 3 µm. ii: Magnified view of tunnel-nucleolus contact in boxed area of image (i). Scale bar: 500 nm. Length expansion factor: 4.0. (K) Model of the organization of ribosome biogenesis-associated components near the nuclear tunnel.

At the nuclear tunnel, more organelles and nuclear compartments are engaged. The NE consists of the nuclear membrane, nuclear lamina, and NPCs, and the nuclear tunnel is no exception. We visualized that the nuclear membrane (Figure 3A), lamina (Figure 1P), and NPCs (Figure 3B) continue from the smooth NE to the nuclear tunnels. Due to the high density of NPCs in nuclear tunnels (Figure 3B), we used NPCs to mark nuclear tunnels to locate other organelles (Figures 3E-G). We observed a thin layer of heterochromatin that coats the nuclear tunnel and connects it with the nucleolus (Figure 3E). Pre-ribosomes, labeled by eIF6, are concentrated at the nucleolar rim and form close contact with the NPCs on the nuclear tunnels (Figure 3F). The direct contact between NPCs and pre-ribosomes may accelerate the export of pre-ribosomes, according to recent studies on pre-ribosome maturation at the nucleolar rim and export towards NPCs^45^. However, proximity to NPCs alone is insufficient for pre-ribosome export; pre-ribosomes need to bind exportin1 to pass through NPCs. Imaging exportin1 revealed two distinct groups based on spatial distribution (Figure 3Gi). One group of exportin1 homogeneously diffuses in the nucleoplasm, consistent with the literature^46^. The other group unexpectedly docks at every NPC (Figures 3Gi&ii), concentrating on the nuclear tunnels (Figures 3Gii-iv). These findings demonstrate how exportin1 and NPCs collaborate to efficiently export ribosomal subunits through nuclear tunnels.

On the cytoplasmic side of the nuclear tunnel, ER was labeled with anti-Sec61B antibodies, revealing that ER coats the cytoplasmic side of nuclear tunnels (Figure 3H), consistent with previous studies^38, 47, 48^. Abundant mature ribosomes marked with anti-RPS3 antibodies were also observed in the nuclear tunnels. In the nuclear tunnels, some ribosomes diffuse in the cytosol, while others colocalize with the ER (Figure 3Hii). These ribosomes inside nuclear tunnels may be assembled from pre-ribosomes immediately after export. Surprisingly, mitochondria were also present in nuclear tunnels (Figure 3I). Unlike ER, mitochondria sometimes do not reach the narrow tip of nuclear tunnels due to their larger diameter (∼500nm)^49^. In addition to membrane organelles, cytoskeleton was observed in nuclear invagination with electron microscopy in previous studies ^16^. We observed microtubules (Figure 3J) and F-actin puncta in the nuclear tunnels, but not actin filaments (Figure S7 and Movie S6).

In summary, the nuclear tunnel brings all the necessary components close to the nucleolus for efficient ribosome biogenesis. Our multicolor super-resolution images elucidated the complex spatial relationships among nuclear invaginations, nucleoli, nuclear lamina, NPCs, exportin1, ER, mitochondria, and microtubules. Figure 3K illustrates these intricate organelle-organelle interactions around the nuclear tunnel.

#### Nuclear tunnels attenuate heterochromatin but enrich NPCs

The next question is whether nuclear tunnels alter the distribution of chromatin or organelles. The answer is clearly yes for the nucleolus, as shown in Figures 3A-C. Now, let’s take a closer look at chromatin. It is known that the majority of heterochromatin in a cell is located at the periphery of the nucleolus and the nuclear lamina, known as nucleolus-associated domains (NADs) and lamina-associated domains (LADs) (Figure 4Ai)^50^. Our new finding is that NADs and LADs dramatically decrease at the contact site between the nucleolus and the nuclear tunnel (Figure 4Bii, arrow), in contrast to the thick NADs and LADs at the contact site between the nucleolus and the flat NE (Figure 4Aii, arrow). We quantified the heterochromatin thickness at the tunnel-nucleolus contact sites and flat-nucleolus contact sites. The statistics show that the heterochromatin domain between a nucleolus and a nuclear tunnel is on average 3 times thinner than the heterochromatin between a nucleolus and a dent or flat NE (Figure 4C). Interestingly, the overall NADs are significantly decreased around the whole nucleolus if it is contacted by a tunnel, allowing rDNAs in this region to be actively transcribed. This attenuation of heterochromatin reveals a possible mechanism on how nuclear tunnels upregulate nucleolar activity. It is known that lack of NADs alters nucleolar structure^51^, and increases ribosome biogenesis^52^.

**Figure 4.**
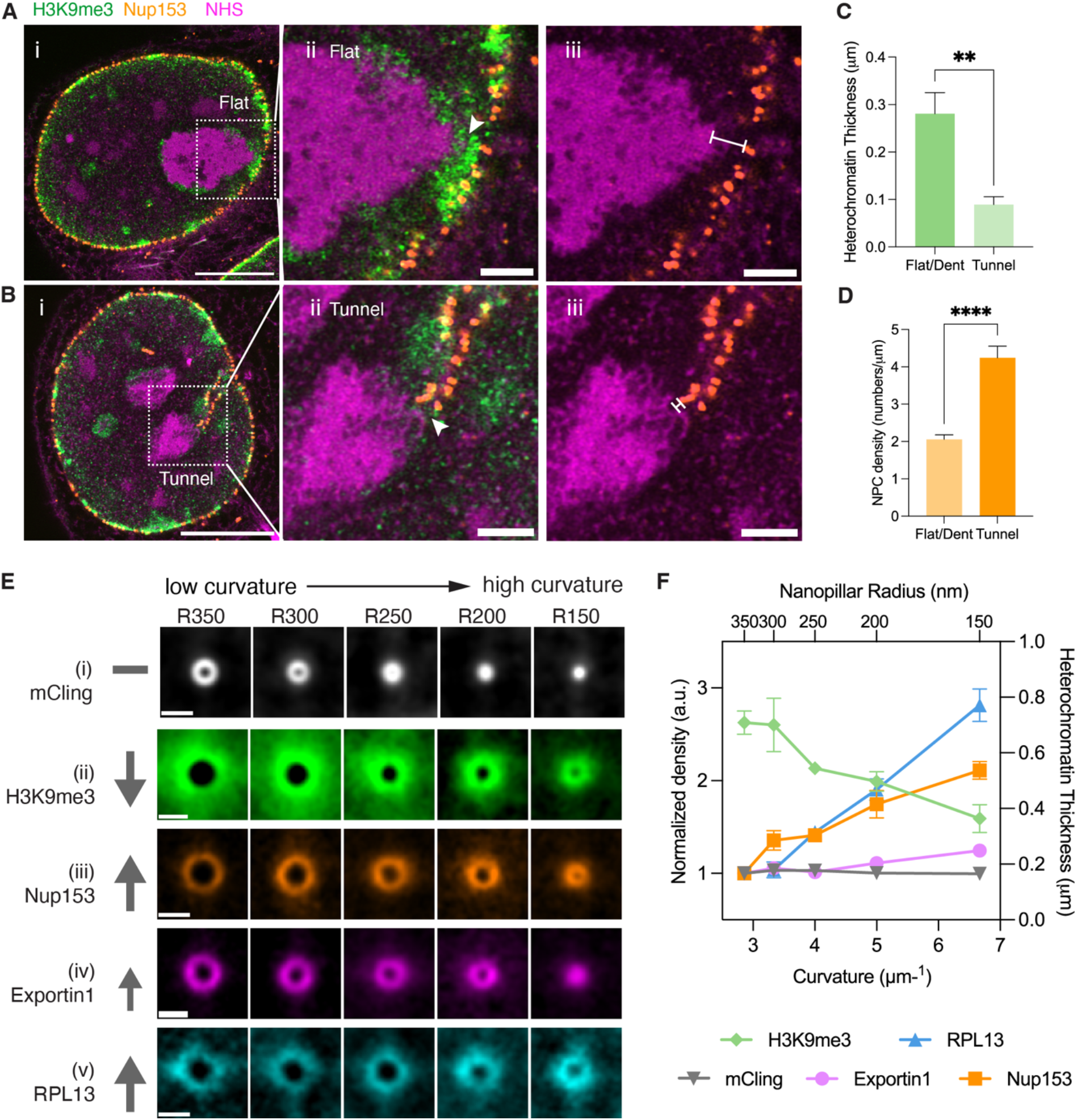
Nuclear Tunnels Reduce Heterochromatin and Enrich Nuclear Pore Complexes. (A) i: Expansion microscopy image of heterochromatin and NPCs at the flat NE-nucleolus contact in an MCF-10A cell. The cell was stained with anti-H3K9me3 antibodies (green), anti-Nup153 (orange), and NHS ester (magenta). Scale bar: 5 µm in pre-expansion unit. ii and iii: Magnified views of NE-nucleolus contact in boxed area of image (i). The arrowhead points to the contact site. The line with flat ends marks the thickness of the heterochromatin between NE and the nucleolus. Scale bar: 1 µm in pre-expansion unit. Length expansion factor: 3.9. (B) i: Expansion microscopy image of heterochromatin and NPCs at the tunnel-nucleolus contact in an MCF-10A cell. The cell was stained with anti-H3K9me3 antibodies (green), anti-Nup153 (orange), and NHS ester (magenta). Scale bar: 5 µm in pre-expansion unit. ii and iii: Magnified views of NE-nucleolus contact in boxed area of image (i). The arrowhead points to the contact site. The line with flat ends marks the thickness of the heterochromatin between NE and the nucleolus. Scale bar: 1 µm in pre-expansion unit. Length expansion factor: 3.9. (C) Barchart of heterochromatin thickness at the flat/dent-nucleolus contact and tunnel-nucleolus contact in MCF-10A cells. Each bar represents the mean ± standard error of more than 15 contacts from 3 independent experiments. ** indicates p<0.01 by unpaired t test. (D) Barchart of NPC density at the flat/dent-nucleolus contact and tunnel-nucleolus contact in MCF-10A cells. Each bar represents the mean ± standard error of more than 15 contacts from 3 independent experiments. **** indicates p<0.0001 by unpaired t test. (E) Images of the cross-sections of nanopillar-induced nuclear invaginations in MCF-10A cells stained with mCling (i), H3K9me3 (ii), Nup153 (iii), Exportin1 (iv), RPL13 (v), respectively. Each image was averaged from more than 250 nanopillar-induced nuclear invaginations. The arrows show the trends in the marked targets as the nanopillar radii decrease from 350 nm to 150 nm. Scale bar: 1 µm. (F) Scatter plot of density of targeted molecules or chromatin thickness at nuclear invaginations induced by nanopillars with radii from 350 nm to 150 nm, which were measured from the images in (E). Each plot represents a mean ± standard error of more than 250 nanopillars from 3 independent experiments. The density or thickness corresponding to each radius was normalized to those corresponding to the radius of 350 nm. All images in this figure were taken on an Airyscan microscope with a 63x objective.

In contrast with the attenuation of heterochromatin, NPCs are enriched on nuclear tunnels (Figure 4Biii). The density of NPCs on the nuclear tunnels is 2 folds higher than that on nuclear dents or flat NE (Figure 4D). The enriched NPCs at the tunnels promote the export rate of pre-ribosomes. The heterochromatin may also serve as a diffusion barrier for pre-ribosomes since its condensed feature, in addition to regulating the transcription of rDNAs. The attenuation of heterochromatin speeds up the diffusion of pre-ribosomes. Therefore, the combination of heterochromatin attenuation and NPC enrichment accelerates the export of pre-ribosomes in nuclear tunnels. The altered distribution of heterochromatin and NPCs at nuclear tunnels motivated us to quantitatively examine the impact of invagination curvature on the organization of chromatin and NPCs.

#### Curvature-dependent arrangement of heterochromatin and NPCs

We used nanopillars to investigate whether the arrangement of heterochromatin and NPCs depends on the curvature of their host nuclear invaginations. We cultured MCF-10A cells on the same nanopillar substrates previously used to study the curvature dependency of nucleolar activity (Figure S5). During cell culture, these nanopillars, with varying radii, generated nuclear invaginations of distinct curvatures in a controllable manner.

Figure 4Ei shows cross-sections of nanopillar-induced nuclear invaginations, demonstrating the controlled size in high-resolution. The nuclear invaginations were stained with a lipid dye, mCling. These images indicate an uniform nuclear membrane density across different curvatures, serving as a surface area control for changes in chromatin, NPCs, and exportin1. Notably, heterochromatin thickness and abundance decreased with ascending curvature (Figure 4Eii). Conversely, the densities of NPCs and exportin1 increased with higher curvature (Figures 4Eiii and 4Eiv).

We employed the fluorescence intensity of ribosomal protein RPL13 to report consequences in ribosome biogenesis. RPL13 had increased levels with higher curvature at the NE-nucleolus contact (Figure 4Ev). Figure 4F summarizes the trends in the abundance of heterochromatin, NPCs, exportin1, and ribosomes with increasing invagination curvature. From a curvature range of 2.5 to 6.7 μm⁻¹, the density of NPCs doubled, exportin 1 increased by 20% while the thickness of heterochromatin significantly decreased by 50% (Figure 4F). As an outcome in ribosome biogenesis, RPL13 nearly tripled. These findings suggest that heterochromatin and NPCs respond to the curvature of the nuclear envelope and alter ribosome biogenesis.

#### Model of biophysical process of ribosome biogenesis

With experimental results, we have demonstrated that the arrangement of chromatin and organelles highly depends on the curvature of the nuclear invagination. Here, we use computational simulation to provide a biophysical interpretation of the curvature-dependent regulation of ribosome biogenesis.

The model describes a two-dimensional side view of the nucleus, with pre-ribosomes releasing from the nucleolar surface, diffusing in chromatin and nucleosol, binding to exportin1, and exporting through NE via NPCs, shown in Figure 5A. The concentration of released pre-ribosomes, *p*, is controlled by the nucleolar production rate. The heterochromatin, with thickness *L* between the nucleolus and the NE, serves as a diffusion barrier to pre-ribosomes and slows down their movement. Prior to export, pre-ribosomes must bind to exportin1, so the fraction of pre-ribosomes bound to exportin at NPCs is assumed to be proportional to the exportin1 concentration, *E*. In lieu of incorporating fine-grained details of NPCs, we follow^53, 54^ and model NPC export by a semi-absorbing surface with a reactivity parameter k that encodes NPC density. The parameters *k*, *p*, *L*, and *E* encode the biophysical processes underlying ribosome biogenesis in the nucleus. These parameters all have been measured here as experimental inputs, which vary between different curvatures of nuclear invaginations. The model synthesizes the steady-state flux of ribosomes through the NE corresponding to an invagination curvature (Figure 5B).

**Figure 5.**
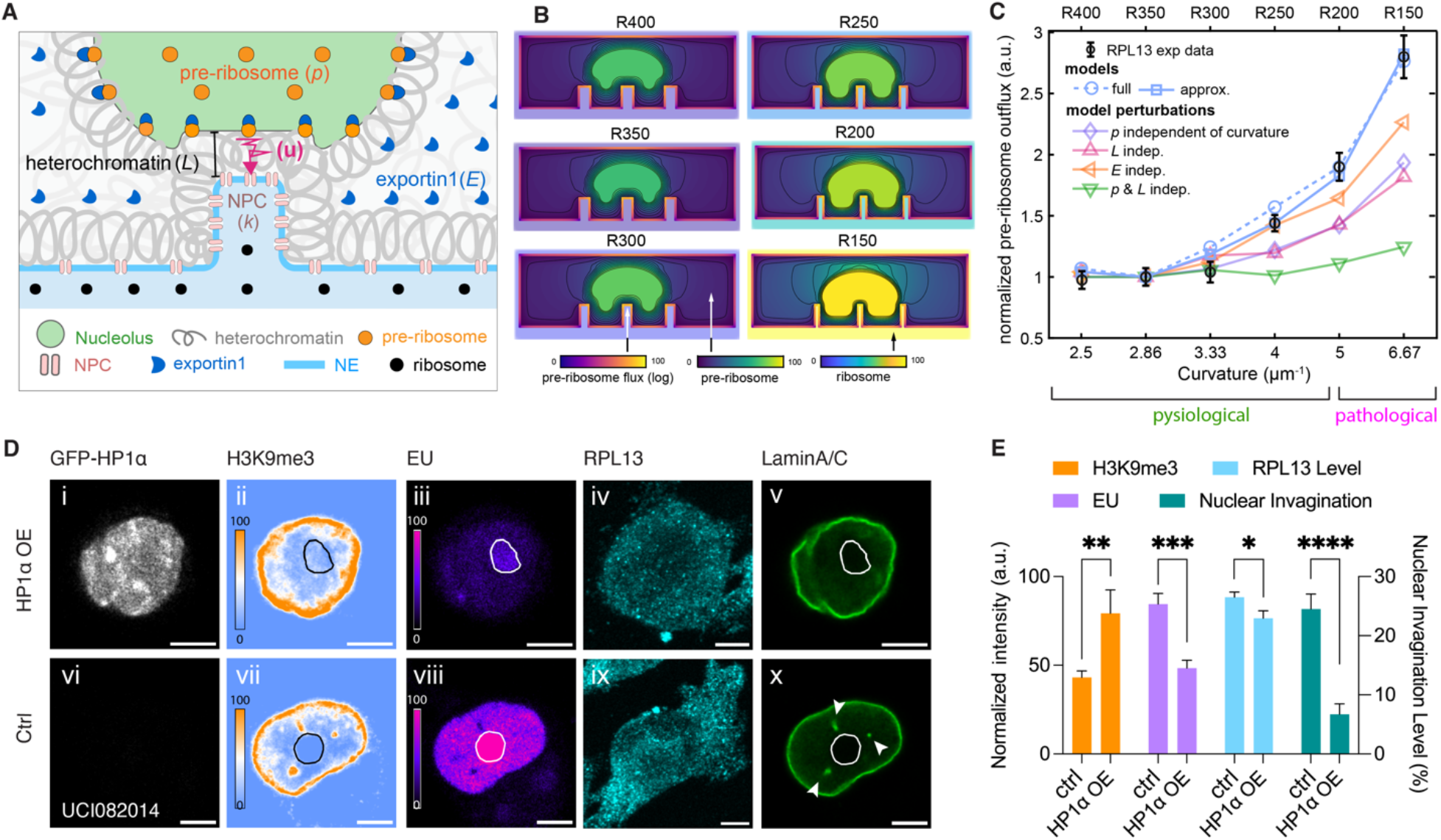
Heterochromatin near NE-nucleolus Contact Plays an Essential Role in Ribosome Biogenesis Regulation. (A) Illustration of the diffusion model of ribosome production in the nucleolus, diffusion transport in the nucleus, and export through NPCs. (B) Simulated 2D distribution of pre-ribosome in the nucleus and ribosome distribution in the cytosol in cells with nuclear invaginations with radii of 400 nm (R400), 350 nm (R350), 300 nm (R300), 250 nm (R250), 200 nm (R200), and 150 nm (R150), respectively. (C) Scatter plots of pre-ribosome outflux in nuclei with different nuclear invagination curvature. All pre-ribosome outflux values are normalized to that from the invagination curvature of 2.5 μm^-1^. The black circles are data points from experimental measurement. The dashed blue curve is simulated by the full 2D diffusion model. The solid blue curve is simulated by the approximate model. The other curves are simulations independent of production rate (purple), independent of heterochromatin thickness (pink), independent of exportin1 (orange), and independent of both production rate and heterochromatin thickness (green), respectively. (D) Airyscan images of UCI082014 cell overexpressing GFP-HP1α (grey) (i-v) and control cells (vi-x) pulse-labeled with EU (magenta hot) for 1 hour and then immunostained with anti-H3K9me3 antibodies (blue-orange), anti-RPL13 antibodies (cyan), and anti-LaminA/C antibodies (green). Nucleoli are outlined with black circles (ii and vii) or white circles (iii, v, viii, and x). Arrows in the image (x) point to nuclear tunnels. Scale bars: 5 µm. (E) Barcharts of the density of H3K9me3, EU, RPL13, and the nuclear invagination levels in HP1α overexpression and control cells. Each bar represents the mean ± standard error of more than 15 cells from 3 independent experiments. * indicates p<0.05, ** indicates p<0.01, *** indicates p<0.001, and **** indicates p<0.0001 by unpaired t test.

We first ask whether the model is sufficient to capture the dependence of ribosome biogenesis on the invagination curvature that was observed in nanopillar experiments (Figure 4C, RPL13 curve). Below a curvature of 5 μm^-1^, the simulated flux remains low within its physiological range. This result agrees with our experimental measurement from cells treated with low-curvature nanopillars (R400, R350, and R300) and measurements in natural nuclear dents (Figure 1U). At the highest curvature of 6.7 μm^-1^, pre-ribosomes outflux increases to pathological level, which is 2.7 times higher than the flux at 2.5 μm^-1^. The agreement between the simulated and experimental data indicates that a diffusion-mediated process is sufficient to explain and interpret the curvature-dependent regulation of ribosome biogenesis (Figure 5C). For deeper insight, we turn to an approximation of the full model that permits an interpretable analytical answer for the output flux. We reduced the full model to the passage directly between the nucleolus to the NE, and from this approximation, compute the mathematical expression for the output flux (supplementary information). The expression for output flux reveals the insight that output is a fine balance between diffusion-limited and NPC-limited transport. Simulations of the approximate model show that total export is dominated by the region where the nucleolus is in closest proximity to the NE. Remarkably, this reduced model still reflects the RPL13 measurements faithfully.

With the model validated, we used it to dissect how the intertwining biophysical factors shape the overall export rate. It is challenging to decouple the regulators experimentally, yet, straightforward for the model. To understand quantitively how each biophysical factor contributes to the curvature-dependent increase in flux, we singled out the impact by setting its corresponding parameter with a constant across different curvatures. For instance, to investigate the nucleolar contribution, we set the production rate of pre-ribosomes (*p*) as a curvature-independent with the value of the control and kept *L* and *E* varying with experimentally measured curvature-dependence. The curve of the flux maintains the climbing trend but with a lower slope (Figure 5C, purple curve). As a result, pre-ribosome outflux can be increased by 1.8 times at the highest curvature, which is not as significant as the 2.7 times increase in the original model and experimental results. Similarly, by setting the heterochromatin thickness (*L*) or exportin1 constant at the control value, the model predicted the manipulated curves of pre-ribosome outflux with slower slopes (Figure 5C, pink and orange curves). The larger the difference between the pre-ribosome flux of the manipulated curve and the best-fitting curve, the greater the influence of the investigated organelle. Therefore, we conclude that both nucleolus and heterochromatin are the major contributors to the regulation of ribosome biogenesis, while exportin has a substantial yet smaller contribution. Since the abundance of heterochromatin affects both nucleolar activity and pre-ribosome diffusion, we also simulated a curve with constant ribosome production (*p*) and constant heterochromatin thickness (*L*) at their control values. Consequently, the curve of pre-ribosome flux flats out (Figure 5C, green curve). These modeling results emphasize the importance of heterochromatin abundance in regulating ribosome biogenesis.

#### Heterochromatin enrichment reduces ribosome biogenesis and nuclear invaginations

Our model predicted that increasing heterochromatin would reduce ribosome biogenesis. Therefore, switching towards a heterochromatin stage might be sufficient to repress the overactive ribosome biogenesis observed in cancer cells.

We experimentally validated this theoretical prediction in the triple negative breast cancer UCI082014 cells (Figure 5D). HP1α, a heterochromatin protein known to promote heterochromatin^55^, was overexpressed in UCI082014 cells (Figure 5Di). We immunostained the heterochromatin marker H3K9me3 in both HP1α-overexpressing cells and control UCI082014 cells. The overexpression resulted in an increased amount of heterochromatin at both the nuclear periphery and nucleolar periphery (Figure 5Dii), compared to the control group (Figure 5Dvii). To assess the impact on ribosome biogenesis, we immunostained ribosomes with anti-RPL13 antibodies and labeled RNAs with EU. The EU signal in the nucleoli reflects the amount of rRNA. Figure 5Diii shows that the amount of newly synthesized rRNA was significantly decreased in cells overexpressing HP1α, compared to the control (Figure 5Dviii). These results confirmed the model’s prediction that heterochromatin enrichment downregulates ribosome biogenesis. As a consequence of the inactive rRNA synthesis, ribosome levels in the cytosol of HP1α-overexpressing cells were also significantly reduced (Figure 5Div) compared to the control group (Figure 5Dix). These results show that heterochromatin enrichment represses the initial step of ribosome biogenesis: rDNA transcription and rRNA synthesis.

Is heterochromatin enrichment an independent rescue approach, separate from our structural approach that converts nuclear tunnels to dents using nanopillars? The answer is no; they are interdependent. We found that cells with enriched heterochromatin lost nuclear tunnels, which are associated with the downregulation of ribosome biogenesis. Figure 5Dx displays the typical morphology of UCI082014 cells, which have multiple high-curvature nuclear tunnels (indicated by white arrowheads). These nuclear tunnels contacted the nucleolus in the center. In contrast, nuclei with overexpressed HP1α reduced high-curvature invaginations (Figure 5Dv). The results indicate that heterochromatin enrichment can remove high-curvature nuclear invaginations and simultaneously suppress ribosome biogenesis, a finding that is statistically rigorous (Figure 5E). This finding evoked the final question of this work: what causes nuclear invaginations?

### Direct causes of nuclear deformation

Nuclear morphology is determined by the force balance from both sides of the NE and the stiffness of the nuclear lamina. A recent study on heterochromatin-driven nuclear softening revealed that the loss of heterochromatin as a rapid response to cause high-curvature nuclear invaginations^56^. Their results also showed that once the heterochromatin is rebuilt over time, the nuclear invaginations disappear. This report helps interpret why we observed the loss of high-curvature nuclear invaginations in cells with enhanced heterochromatin (Figures 5Dv & 5E).

Now, we turn to another determinant of nuclear morphology: the nuclear lamina. It is well known that when the nuclear lamina is stiff, it is more resilient to forces from the chromatin or cytoplasm. The nuclear lamina, a dense filament network beneath the inner nuclear membrane, consists of two types of lamin isoforms: A-type lamin and B-type lamin. Extensive studies have shown that A-type lamins are associated with nuclear stiffness and deformability^3,27, 57^. Therefore, we questioned if lamin isoforms are involved in the formation of nuclear invaginations.

We first examined the distribution of the two types of lamina isoforms at nuclear tunnels and the flat NE. We co-stained lamin A/C and lamin B1 in the breast cancer cell line MDA-MB-231, which contains abundant nuclear invaginations. Our super-resolution images in Figures 6A & 6B show that lamin B1 preferentially distributes at the nuclear tunnel, while lamin A/C has no preference between the nuclear tunnels and flat NE. We examined PDX-based breast cancer tumors and observed the same trends in the distribution of lamin isoforms (Figures 6C & 6D). We suspect the B-type to A-type lamin ratio correlates with the curvature of nuclear invaginations. To validate the correlation, we cultured cells on nanopillars with gradient radii and imaged lamin A/C and lamin B1 at the cross-sections of nanopillar-induced nuclear invaginations. Figures 6E & 6F show that the lamin B1 density increases with ascending nanopillar curvature, while lamin A/C density remains roughly consistent. The lamin B1 to lamin A/C ratio positively correlates with the curvature of nuclear invaginations (Figure 6G). Thus, we hypothesized that high lamin B1 to lamin A/C ratios can cause nuclear invaginations.

**Figure 6.**
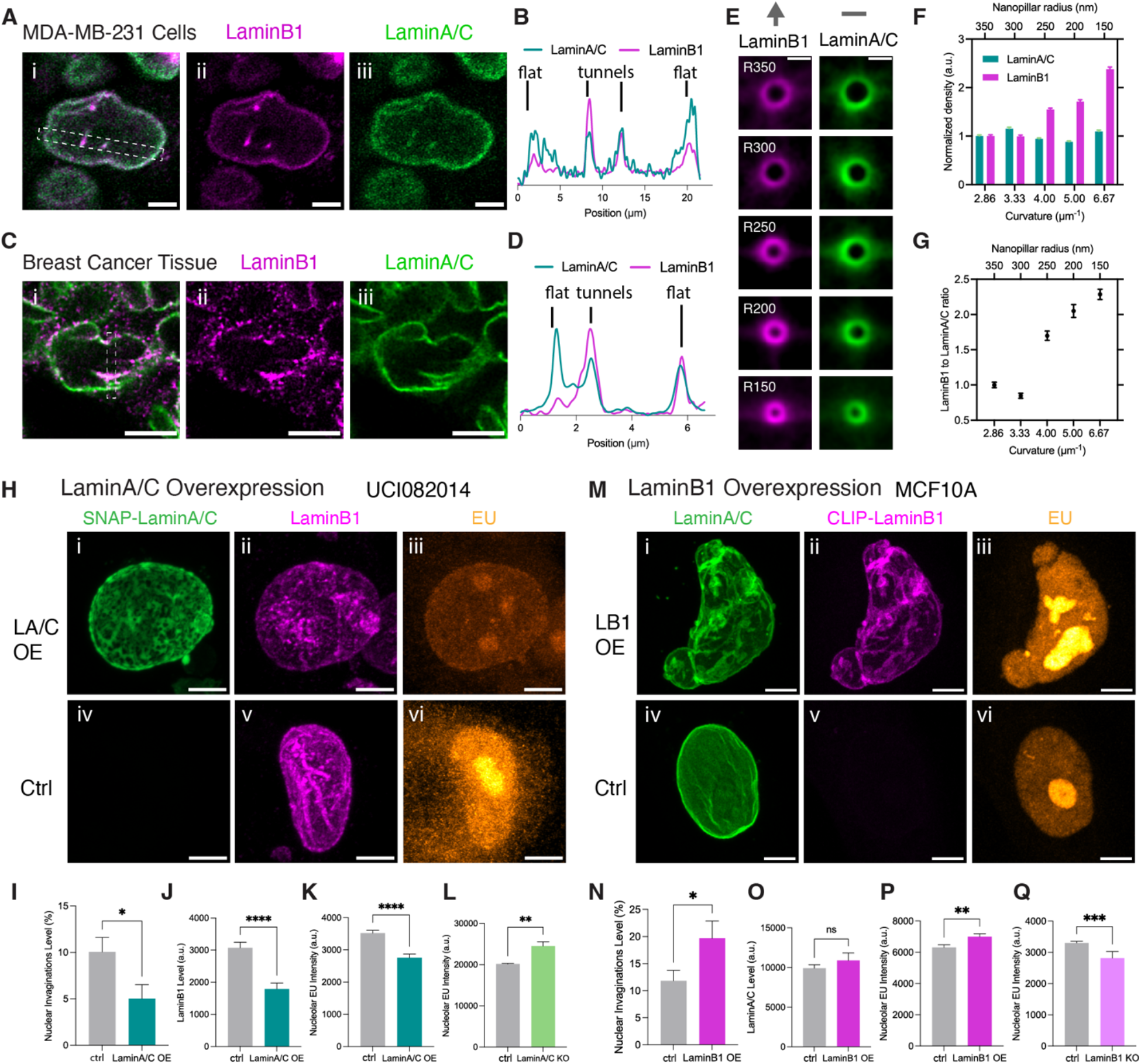
High B-type to A-type Lamin Ratios Cause Nuclear Invagination and Elevate Ribosome Biogenesis. (A) Images of MDA-MB-231 cells immunostained with anti-LaminB1 (magenta) and anti-LaminA/C (green) antibodies. Scale bar: 5 µm. (B) Transverse intensity profile of LaminB1 and LaminA/C in the boxed area of the image (A). (C) Images of breast cancer patient-derived xenograft immunostained with anti-LaminB1 (magenta) and anti-LaminA/C (green) antibodies. Scale bar: 5 µm. (D) Transverse intensity profile of LaminB1 and LaminA/C in the boxed area of the image (C). (E) Average images of LaminB1 (magenta) and LaminA/C (green) at the cross-sections of nuclear tunnels generated by nanopillars with gradient radii from 350 nm to 150 nm in MCF-10A cells. Each image was averaged from more than 250 nuclear invaginations. The arrow on top of LaminB1 images shows the increase of LaminaB1 density as the nanopillar radii decrease from 350 nm to 150 nm. The short line above LaminA/C images indicates the unchanged level of LaminA/C as the invagination radii ascend. Scale bar: 1 µm. (F) Barchart of LaminB1 and LaminA/C density at the nuclear invaginations generated by nanopillars with gradient radii from 350 nm to 150 nm in MCF-10A cells. Each bar represents the mean ± standard error of more than 250 nanopillars from 3 independent experiments. The density of the lamin proteins at each radius is normalized to that at the radius of 350 nm. (G) Scatter plot of LaminB1 to LaminA/C ratios at nuclear invaginations with different curvatures. The ratios are calculated from the data from (F). (H) Images of UCI082014 cell overexpressing snap-LaminA/C (green) (i-iii) and control cells (iv-vi). The cells were pulse-labeled with EU (orange) for 1 hour and then immunostained with anti-LaminB1 antibodies (magenta). Scale bar: 5 µm. (I-K) Barcharts of nuclear invagination level (I), LaminB1 (J), and EU intensity (K) in cells overexpressing LaminA/C and control cells. Each bar represents the mean ± standard error of more than 50 cells from 3 independent experiments. * indicates p<0.05 and **** indicates p<0.0001 by unpaired t test. (L) Barchart of EU intensity in LaminA/C knockout and control cells. Each bar represents the mean ± standard error of more than 20 cells from 3 independent experiments. ** indicates p<0.01 by unpaired t test. (M) Images of MCF-10A cells overexpressing clip-LaminB1 (magenta) (i-iii) and control cells (iv-vi). The cells were pulse-labeled with EU (orange) for 1 hour and then immunostained with anti-LaminA/C antibodies (green). Scale bar: 5 µm. (N-P) Barcharts of nuclear invagination level (N), LaminA/C (O), and EU intensity (P) in cells overexpressing B1 and control cells. Each bar represents the mean ± standard error of more than 50 cells from 3 independent experiments. * indicates p<0.05, ** indicates p<0.01 and ns indicates p>0.05 by unpaired t test. (Q) Barchart of EU intensity in LaminB1 knockout and control cells. Each bar represents the mean ± standard error of more than 50 cells from 3 independent experiments. *** indicates p<0.001 by unpaired t test. All images in this figure were taken on an Airyscan microscope with a 63x objective.

To test this hypothesis, we altered the expression levels of lamin isoforms using overexpression and knockout techniques. First, we overexpressed SNAP-lamin A/C in UCI082014 breast cancer cells, which initially have many high-curvature nuclear invaginations. Nuclei with overexpressed lamin A/C lost high curvature nuclear invaginations and became rounder with fewer invaginations (Figures 6Hi & 6I) compared to the control cells (Figure 6Hv). Consequently, less rRNA was synthesized by the nucleolus (Figures 6Hiii & 6K). The downregulation of nucleolar ribosome biogenesis confirmed our findings of curvature-dependent regulation of ribosome biogenesis (Figures 1A, 2J, and 2K). Surprisingly, lamin A/C overexpression also caused a substantial reduction of lamin B1 (Figures 6Hii & 6J). This result indicates that lamin A/C can regulate lamin B1 levels, further lowering the lamin B1 to lamin A/C ratio. Second, we knocked out lamin A/C in MCF-10A cells. The MCF-10A cells rarely have nuclear invaginations. As predicted by the hypothesis, the lamin A/C knockout led to more nuclear invaginations and increased rRNA synthesis in nucleoli (Figure 6L). Third, we overexpressed lamin B1 in MCF-10A cells, and the nuclear invaginations exaggeratedly increased (Figures 6Mii & 6N). As expected, more rRNA was synthesized by the nucleoli in the lamin B1 overexpressing cells (Figures 6Miii & 6P). Although the overall lamin A/C level remained unchanged (Figure 6O), the lamin B1 to lamin A/C ratio increased due to the overexpression of lamin B1. Finally, lamin B1 knockout lowered the lamin B1 to lamin A/C ratio and simultaneously deactivated rRNA synthesis in UCI082014 cells (Figure 6Q). Altogether, our results proved that high lamin B1 to lamin A/C ratios can cause nuclear invaginations. Tuning down the lamin B1 to lamin A/C ratio can effectively reduce ribosome biogenesis.

## Discussion

We position our new findings within the current understanding of nuclear deformation to discuss their significance and connections with other significant discoveries.

Since the 1950s, nuclear invaginations have been observed with the electron microscopy^58^. Bourgeois *et al*. further confirmed the surprising contact between a nuclear tunnel and a nucleolus^13^. However, capturing the NE-nucleolus contact sites is statistically challenging for electron microscopy because of the low chance of having the perfect sectioning at the contact sites. 2D electron microscopy images of the wrong z plane will miss the contact sites or overestimate the distance between the nuclear invagination and the nucleolus. Therefore, 3D super-resolution imaging is needed to reliably display the spatial relationship between the nuclear invaginations and the nucleoli. With advancements such as expansion microscopy, light microscopes like Airyscan now achieve effective resolutions of 30 nm, comparable to SEM but with 3D imaging capability and high speed. Using LR-ExM Airyscan, we observed the same nuclear tunnel-nucleolus contact and quickly examined hundreds of nuclei. Now we can answer the fundamental question: how often do nuclear invaginations contact nucleoli? Our 3D images indicate that most nucleoli are contacted by the NE across various cell lines and tissues, with suspended nucleoli being a minority. Furthermore, the stability of NE-nucleolus contact during the cell cycle was investigated. Our live-cell imaging shows that the nuclear tunnel-nucleolus contact lasts for hours and persists throughout the interphase (Movie S3), demonstrating the statistical significance of these contacts.

Despite the decades since the first observation of nuclear tunnel-nucleolus contact, its functional consequences remain unclear. This knowledge gap hampers our understanding of critical cell types, such as stem cells and neutrophils, which have abundant nuclear invaginations. It also impedes our understanding of diseases characterized by severe nuclear invaginations, including cancers, progeria neurodegeneration, and infections. Our research uncovers a significant and surprising function of nuclear invaginations: regulating ribosome biogenesis. Given the critical role of ribosome biogenesis in protein synthesis, cell growth, proliferation, differentiation, and apoptosis, its deregulation can lead to various diseases. The link between nuclear invagination and nucleolar activity helps explain why ribosome biogenesis is upregulated in many diseases, such as breast cancers, progeria, and normal aging^10, 11, 59^. We observed that rRNA and polI levels increase in nucleoli contacted by high-curvature nuclear invaginations, directly evidencing this upregulation.

Our most striking finding is that the regulation of nucleolar activity is precisely localized. Each nucleolus can be individually activated by contact with a nuclear invagination. The curvature of the invagination determines the rate of rRNA synthesis and pre-ribosome assembly through physical contact. The higher the curvature, the higher the rate of ribosome biogenesis in the contacted nucleolus. In a natural nucleus (Figure 1P) or a half-nucleus deformed by nanopillars (Figure 2H), only nucleoli contacted by nuclear tunnels show increased ribosome biogenesis. This regulatory mechanism is distinct from canonical regulation pathways like the mTOR1 pathway, which involves the nuclear translocation of TIF-IA, a transcription factor critical for polI-mediated rRNA synthesis^60^. Our imaging showed phosphorylated mTOR distributed in the nucleus without spatial preference at individual nucleoli or nuclear tunnels (Figure S8A), although its total nuclear level can be increased by high-curvature nanopillars (Figure S8B). Therefore, we conclude that local contact between nuclear invaginations and nucleoli represents a novel regulatory mechanism distinct from signaling pathways.

It is common to see local arrangement of proteins or organelles by membrane curvature in cells. Curvature, a mathematical concept quantifying edge deviation from a straight line, dynamically affects the distribution of lipids and membrane proteins^61^. Membrane curvature has been extensively demonstrated as a mechanism to precisely control biochemical reaction rates in cells ^62^. For instance, at the plasma membrane, the maturation step of endocytosis depends on the curvature of endocytic sites^63, 64^. Plasma membrane curvature also determines the wave propagation speed in immune cells^65^. In the nucleus, the NE curvature facilitates nuclear pore complex assembly^61, 66^. Compared to other membranous structures, the role of membrane curvature in the nucleus is less understood. Our study, using nanopillars high enough to deform nuclei, reveals a linear positive correlation between the rate of ribosome biogenesis and the nuclear invagination curvature. This curvature-dependent regulatory mechanism explains why in cancer and progeria cells with abundant nuclear tunnels and folds have overactive ribosome biogenesis. The quantitative measurement of curvature-dependency also provides a curvature threshold to determine whether a nuclear deformation is benign and pathological for ribosome biogenesis. This threshold is about 5 μm⁻¹. Nuclear invaginations with a curvature bigger than 5 μm⁻¹ will promote ribosome biogenesis in the contacted nucleolus to a pathological level.

While curvature sensing primarily deals with the shape and morphology of cellular membranes, mechanosensing focuses on the forces exerted on cells. Both processes often overlap, as changes in membrane curvature can affect mechanical forces and vice versa. However, the nucleolus is not primarily known for being curvature-sensitive or mechanosensitive. We suggest that the nuclear invagination function as an intermediate regulator which impacts the nucleolus through interactions with curvature-sensitive or mechanosensitive structures and organelles. Our data show that at least three mechanosensitive structures relay between the invaginated NE and the nucleolus, which are the nuclear lamina, NPCs, and heterochromatin (Figure 4 and 6). The nuclear lamina and NPCs are components of NE, and the heterochromatin domains are anchored to the NE lamina by interacting with nuclear lamina-binding proteins, such as lap2β, emerin, and lamin B receptor ^67–69^. Thus, it is not surprising to see them following the nuclear invaginations. The surprising finding here is their arrangement at the nuclear invagination is distinct from the smooth NE, and their abundance at the invaginations depends on the curvature. This curvature-dependent distribution of heterochromatin and NPCs can cause curvature-dependent ribosome biogenesis in the nearby nucleolus. It is because that heterochromatin regulates the rDNA transcription and NPCs facilitate the export of pre-ribosome subunits^44, 52^. This new regulatory mechanism of ribosome biogenesis advanced our understanding of the function of nuclear invaginations.

As discussed, nuclear invaginations may function as an intermediate regulator of ribosome biogenesis in the structure-function pathway. In addition to the downstream regulator, such as heterochromatin, any causes for nuclear invaginations could be upstream regulators of ribosome biogenesis. In the scope of this study, we only investigated several direct causes of nuclear invagination, such as B-type to A-type lamin ratio, heterochromatin, and microtubules. Several more direct and indirect causes of nuclear deformation have been reported, such as p53, Hippo, and Akt signaling pathways^70–72^. These signaling pathways will very likely affect ribosome biogenesis through the nuclear invagination structure, which requires future investigation.

## Conclusions and future directions

This work has reshaped our understanding of nuclear invaginations, transitioning them from previously vague descriptors of nuclear deformation to well-defined structures with a pivotal role in regulating ribosome biogenesis. Our findings address three significant questions in the nuclear invagination: what its relationship with ribosome biogenesis is, how it works, and why it forms.

Firstly, we unveil a novel function of nuclear invaginations: the upregulation of ribosome biogenesis. Our experiments reveal that this regulation is contingent upon the curvature of the nuclear invagination, as not all nuclear invaginations affect ribosome biogenesis. Only those exhibiting high curvature have the potential to enhance nucleolar activity through physically contacting the nucleolus. While not necessarily the initial cause, the nuclear invagination serves as an intermediate regulator of ribosome biogenesis. Remarkably, altering the curvature of nuclear invaginations through nanopillars effectively reduces pathological levels of ribosome biogenesis to physiological levels. This result introduces a novel structural approach for mitigating inflated ribosome biogenesis without genetic editing or pharmaceutical intervention.

Second, we elucidate the cellular mechanism and explored the causes of nuclear invaginations. Our observation uncovered that the nuclear invagination organizes various cellular components in its proximity—such as the nucleolus, heterochromatin, nuclear lamina, NPCs, exportin1, ER, mitochondria, and microtubules—within a confined subcellular compartment. What’s important is that the arrangement of all these components in the nuclear invaginations is dependent on the invagination curvature. High-curvature nuclear invaginations attenuate heterochromatin, and enrich NPCs and exportins, all of which promote ribosome biogenesis. This implies that nuclear invagination put together teamwork of organelles and complexes to drive ribosome biogenesis. While our experiments showed orchestrated contributions to ribosome biogenesis, our biophysical modeling separated the contributions from each component. The model highlights the significant role of heterochromatin loss on high-curvature nuclear invaginations as a major promoter of ribosome biogenesis—a finding we experimentally validated.

Finally, we conducted a preliminary exploration of the causes of nuclear invaginations. We found that softer chromatin with reduced heterochromatin or a less rigid nuclear lamina increases the propensity for nuclear invagination. A low ratio of A-type to B-type lamin isoforms results in a softer NE, thereby favoring high-curvature invaginations. Notably, we observed heterogeneous ratios within an individual nucleus, with lower ratios of A-type to B-type lamins correlating with high-curvature nuclear invaginations. We anticipate that future investigations into upstream factors, such as LMNA mutations and aberrant signaling pathways, will further elucidate the intricate etiology of nuclear invaginations.

Our study owes much to emerging biotechnologies, particularly expansion microscopy and nanopillar materials. LR-ExM provided the resolution, imaging depth, multiplexing capability, and throughput necessary for the comprehensive analysis of nuclear invaginations’ structure, function, and mechanisms. The application of nanopillars demonstrated a high-resolution control of the nanoscale nuclear invagination and led to the discovery of its curvature-dependent regulation of ribosome biogenesis. Excitingly, nanopillars also showcased the potential to modulate cellular functions—such as ribosome biogenesis—by altering the underlying structure, ushering in a paradigm where structure-function relationships can be directly leveraged for therapeutic intervention.

While our work represents a significant step towards unraveling the functions of nuclear invaginations, it merely scratches the surface. We acknowledge that nuclear invaginations may interact with other organelles beyond those associated with ribosome biogenesis. A comprehensive exploration of these interactions will promise to unveil new functions of nuclear invaginations. Moreover, our study did not delve into the molecular mechanisms underpinning the structure-function relationship, leaving room for proteomic and molecular biological studies to elucidate the molecular drivers of curvature-dependent regulation in the future.

## Supplemental information

Document S1. Figures S1-S10

Movie S1. 3D stack of Airyscan expansion microscopy images of UCI082014 cell from the bottom to the top. The cell was stained with total lipid (magenta) and total protein (cyan) dyes. The length expansion factor: 3.9. Scale bar: 5 µm in pre-expansion unit. Related to Figure 1.

Movie S2. 3D stack of Airyscan expansion microscopy images of UCI082014 cell from the bottom to the top. The cell was stained with total lipid (grey) and total protein (red) dyes. The length expansion factor: 3.8. Scale bar: 5 µm in pre-expansion unit. Related to Figure 1.

Movie S3. Airyscan live cell imaging of nuclear tunnel-nucleolus contact. HEK293T cell was CRISPR knocked in split-mNeongreen UBTF (red hot) and stained with ERtracker (cyan). Imaging interval: 2 min/frame. Scale bar: 5 µm. Related to Figure 1.

Movie S4. Confocal live cell imaging of UCI082014 cells stained with ERtracker (cyan). Red arrowhead indicates cell mitosis. Imaging interval: 5 min/frame. Scale bar: 10 µm. Related to Figure 1.

Movie S5. Airyscan live cell imaging of nuclear dent-nucleolus contact. HEK293T cell was CRISPR knocked in split-mNeongreen UBTF (red hot) and stained with ERtracker (cyan). Imaging interval: 2 min/frame. Scale bar: 5 µm. Related to Figure 1.

Movie S6. 3D stack of Airyscan expansion microscopy images of UCI082014 cell from the bottom to the top. The cell was stained with total lipid (magenta), total protein (grey) dyes, phalloidin-fluorescein, and anti-fluorescein antibody (green). The length expansion factor: 3.8. Scale bar: 5 µm in pre-expansion unit. White arrowheads indicate the puncta-like F-actin. Related to Figure 3.

## Star Methods

### Key Resources Table

| Materials and Reagents | Source | Identifier |
| --- | --- | --- |
| **Antibody conjugation or whole protein staining** |  |  |
| (+)-Biotin N-hydroxysuccinimide ester | Sigma | H1759 |
| Digoxigenin N-hydroxysuccinimide ester | Enzo | ENZ-45022-0001 |
| Methacrylic acid N-hydroxysuccinimide ester | Sigma | 730300 |
| Alexa Fluor™ 405 NHS Ester (Succinimidyl Ester) | Invitrogen | A30000 |
| AF 488 NHS ester | Lumiprobe | 21820 |
| Alexa Fluor™ 568 NHS Ester (Succinimidyl Ester) | Invitrogen | A20003 |
| Alexa Fluor™ 647 NHS Ester (Succinimidyl Ester) | Invitrogen | A20006 |
| AF 647 NHS ester | Lumiprobe | 26820 |
| AffiniPure™ Goat Anti-Rabbit IgG (H+L) | Jackson ImmunoResearch | 111-005-144 |
| AffiniPure™ Goat Anti-Mouse IgG (H+L) | Jackson ImmunoResearch | 115-005-166 |
| AffiniPure™ Goat Anti-Rat IgG (H+L) | Jackson ImmunoResearch | 112-005-167 |
| Cytiva NAP-5 columns | Fisher Scientific | 45-000-151 |
| **RNA labeling** |  |  |
| 5-Ethynyl Uridine | Click Chemistry Tools | 1261-10 |
| AZDye 405 Azide | Click Chemistry Tools | CCT-1307 |
| AZDye 568 Azide | Click Chemistry Tools | CCT-1291 |
| Click-iT™ RNA Alexa Fluor™ 594 Imaging Kit | Invitrogen | C10330 |
| Copper(II) sulfate pentahydrate | Sigma | 209198 |
| **Primary antibodies and dyes** |  |  |
| Lamin B1 (D9V6H) Rabbit mAb | Cell Signaling Technology | 13435 |
| Lamin B2 (D8P3U) Rabbit mAb | Cell Signaling Technology | 12255 |
| Lamin A/C (4C11) Mouse mAb | Cell Signaling Technology | 4777 |
| Nucleolin (E5M7K) Mouse mAb | Cell Signaling Technology | 87792 |
| GAPDH (14C10) Rabbit mAb | Cell Signaling Technology | 2118 |
| RPA194 Antibody (C-1) | Santa Cruz Biotechnology | sc-48385 |
| Ribosomal Protein L13 Antibody (SS-09) | Santa Cruz Biotechnology | sc-100829 |
| eIF6 Antibody (A-2) | Santa Cruz Biotechnology | sc-390432 |
| Nup153 Antibody (R4C8) | Santa Cruz Biotechnology | sc-101545 |
| Ribosomal Protein S3 Antibody (C-7) | Santa Cruz Biotechnology | sc-376008 |
| Anti-Nup153 antibody [QE5] | Abcam | ab24700 |
| Anti-Histone H3 (tri methyl K9) antibody | Abcam | ab176916 |
| Sec61b (D5Q1W) Rabbit mAb | Cell Signaling Technology | 14648 |
| Tom20 (D8T4N) Rabbit mAb | Cell Signaling Technology | 42406 |
| Exportin-1/CRM1 (D6V7N) Rabbit mAb | Cell Signaling Technology | 46249 |
| p-mTOR Antibody (59.Ser 2448) | Santa Cruz Biotechnology | sc-293133 |
| Fluorescein Phalloidin | Invitrogen | F432 |
| Fluorescein/Oregon Green Polyclonal Antibody, Alexa Fluor™ 488 | Invitrogen | A11096 |
| SNAP-Cell® Oregon Green® | New England Biolabs | S9104S |
| CLIP-Cell™ TMR-Star | New England Biolabs | S9219S |
| **Whole lipid and protein Expansion microscopy** |  |  |
| SYLGARD™ 184 Silicone Elastomer Kit | Ellsworth | 2137054 |
| 12-mm, no. 1 cover glass | Bellco Glass | 1943-10012A |
| Paraformaldehyde | Electron Microscopy Sciences | 50980488 |
| Aqueous Glutaraldehyde EM Grade 10% | Electron Microscopy Sciences | 16120 |
| Triton X-100 | Sigma | 9002-93-1 |
| bovine serum albumin | Fisher Scientific | BP9703100 |
| mCLING-Atto647N | Synaptic Systems | 710 006AT647N |
| Glycidyl methacrylate | Sigma | 151238 |
| Sodium acrylate, 25 g | Santa Cruz Biotechnology | sc-236893 |
| Acrylamide | Sigma | A9099-100G |
| N,N'-Methylenebisacrylamide | Sigma | M7279-25G |
| N,N,N',N'-Tetramethylethylenediamine | Sigma | T9281-100ML |
| Ammonium persulfate | Sigma | A3678-25G |
| Sodium Dodecyl Sulfate | Fisher Scientific | BP166-500 |
| 1 M Tris HCl Buffer, pH 7.5 | Invitrogen | 15567027 |
| Poly-L-lysine solution, 0.1 % (w/v) in H<sub>2</sub>O | Sigma | P8920-100ML |
| Western blotting |  |  |
| mouse anti-rabbit IgG-HRP | Santa Cruz Biotechnology | sc-2357 |
| m-IgG Fc BP-HRP | Santa Cruz Biotechnology | sc-525409 |
| **Plasmids** |  |  |
| GFP-HP1alpha | Addgene | 17652 |
| pSnap-LMNA | Generated in our own lab |  |
| pClip-LMNB1 | Generated in our own lab |  |
| pRP[CRISPR]-mCherry/Puro-hCas9-U6>hLMNB1[gRNA#276] | Vectorbuilder | VB900129-1184bbw |
| pRP[CRISPR]-mCherry/Puro-hCas9-U6>hLMNB1[gRNA#277] | Vectorbuilder | VB900129-0370xdw |
| pRP[CRISPR]-mCherry/Puro-hCas9-U6>hLMNA[gRNA#9142] | Vectorbuilder | VB900129-1183ber |
| pRP[CRISPR]-mCherry/Puro-hCas9-U6>hLMNA[gRNA#9143] | Vectorbuilder | VB900129-0369qgd |

### Resources Availability

#### Lead contact

Further information and requests for resources can be directed to and will be fulfilled by the Lead Contact, Xiaoyu Shi.

#### Materials availability

Plasmids generated in this study are available by request from the Lead Contact.

#### Data and code availability

All data are available in the main text or the supplementary materials. Codes for nanopillar data analysis are available from the Lead Contact upon request. Codes for ribosome biogenesis modeling are available at GitHub https://github.com/chris-miles/RibosomeDiffusionModel.

### Experimental Model and Subject Details

#### Cell lines

This study utilized MCF-10A, MDA-MB-468, MCF-7, UCI082014, MDA-MB-231, MEF, U2OS, progeria patient-derived HGADFN167 (HGPS, LMNA G608G splice site mutation) and HEK293T (Crispr knock-in with split mNeonGreen-UBTF) cells. All cell lines were maintained in humidified incubator with 5% CO_2_ at 37 °C. MCF-10A was cultured in DMEM/F12 (Gibco^TM^, cat#11320033) supplemented with 5% Horse Serum (Gibco^TM^, cat#16050114), 20 ng/mL Epidermal Growth Factor (PeproTech, cat#AF-100-15), 0.5 µg/mL Hydrocortisone (Sigma, cat# H0888-1G), 100 ng/mL Cholera Toxin (Sigma, cat# C8052-.5MG), 10 µg/mL Insulin (Roche, cat# 11376497001) and 1% penicillin- streptomycin- amphotericin B (Sigma, cat# A5955). MEF and HGADFN167 cells were cultured in DMEM-high Glucose supplemented with GlutaMAX^TM^ (Gibco^TM^, cat#10566024), 15% Fetal Bovine Serum (Gibco^TM^, cat#10082147) and 1% penicillin-streptomycin-amphotericin B. All other cell lines were cultured in DMEM-high Glucose supplemented with GlutaMAX^TM^, 10% Fetal Bovine Serum and 1% penicillin- streptomycin- amphotericin B. To compare the ribosome biogenesis level among different breast cell lines, all breast cell lines including the MCF-10A were cultured overnight in DMEM-high Glucose supplemented with GlutaMAX^TM^, 10% Fetal Bovine Serum and 1% penicillin- streptomycin-amphotericin B before the day of collection or EU pulsed labelling (Figures 1 A-C, Figure S1 and S2). All cell lines were tested mycoplasma-free using MycoStrip™ - Mycoplasma Detection Kit (InvivoGen, cat# rep-mysnc-50). All cells used were < 10 passages from thaw.

HGADFN167 was obtained from the Progeria Research Foundation. HEK293T split mNeongreen- UBTF was obtained from OpenCell (ENSG ID: ENSG00000108312).

#### Patient Derived Xenograft (PDX)

The PDX model used in this study is HCI-002 derived from triple negative breast cancer patient and the primary tumor was fresh frozen until further use.

### Method Details

#### Conjugation of fluorescent secondary antibodies and LR-ExM secondary antibodies

To fulfill multi-color airyscan and expansion microscopy imaging, we conjugated the secondary antibodies with different fluorescent dyes or LR-ExM linkers as described in our previous protocol ^17, 18^. Briefly, secondary antibody of interest (1.25 mg/mL) was mixed and incubated with 100 mM NaHCO_3_ and 2 mg/mL N-hydroxysuccinimide (NHS) ester dye for 15 min at room temperature. For LR-ExM linker conjugation, secondary antibody of interest (1.25 mg/mL) was mixed and incubated with 100 mM NaHCO_3_, 170 µM Digoxigenin N-hydroxysuccinimide ester (or Biotin N-hydroxysuccinimide ester) and 170 µM Methacrylic acid N-hydroxysuccinimide ester for 15 min at room temperature. NAP-5 column was equilibrated with PBS during the antibody mixture reaction. The antibody mixture was then added to the NAP-5 column for purification and collection. Nanodrop (Thermo Fisher, NanoDrop One^c^) was used to measure the conjugated secondary antibody concentration. The concentration of our conjugated antibody was around 0.2-0.3 mg/mL. Details of the materials and reagents used are listed in Key Resources Table.

#### Newly synthesized RNA labeling and immunostaining

Cells were seeded at the density of 0.0125x10^6^/well and cultured in 8 well glass bottom slide (ibidi, cat#80827) overnight before the RNA labeling. To label the newly synthesized RNA, cells were incubated in growth medium with 1 mM 5-Ethynyl Uridine (EU) for 1 hour. Fixation with 4% Paraformaldehyde (PFA) in PBS at room temperature for 10 min, permeation with 0.1% Triton X- 100 in PBS (PBST) for 15 min and blocking with 3% bovine serum albumin (BSA) in PBST for 30 min were then proceeded as standard immunostaining preparation. Before the addition of primary antibodies, azide-dye for the EU fluorescent labeling was prepared in the Click-it reaction cocktails as instructed in the manufacturer’s manual (Click-iT™ RNA Alexa Fluor™ 594 Imaging Kit, Invitrogen™, cat# C10330). We customized the reaction cocktails based on the choice of azide-dye. The Click-it reaction cocktails were consisted of 85.6% reaction buffer provided in the kit, 4mM CuSO_4_, 23 µM Azide-dye, and 10% reaction buffer additive provided in the kit. Incubate the cells with the freshly prepared Click-it reaction cocktails containing the azide-dye for 30 min. Remove the Click-it reaction cocktails and wash the cells with PBS. Primary antibodies (1:200 dilution in 3% BSA in PBST) were later added and incubated with cells overnight at 4°C. After 3 times PBS washing, secondary antibodies conjugated with fluorescent dyes (1:200 dilution in 3% BSA in PBST) were added and incubated with cells for 1 h at room temperature. For staining of snap or clip tag fused on the lamin proteins (Figure 6H and M), we incubated the cells with fluorescent snap or clip tag substrate (3-5 µM in 3% BSA in PBST) together with the secondary antibodies for 1 h at room temperature. After 3 times PBS washing, the cells were proceeded to Airyscan imaging. Details of the reagents used are listed in Key Resources Table.

#### Immunostaining of patient-derived xenograft (PDX)

The PDX tumor was dissected out, chopped to pieces, and freshly frozen in liquid nitrogen before cryosectioning. Tissue-Tek OCT (VWR, cat#25608-930) was used to immerse the PDX tumor, followed by snap frozen on dry ice and stored at −80°C. The OCT-embedded tumor was later proceeded to cryosectioning by vibratome at 20 µm and attached at a positive-charged glass slide. For immunostaining, the tumor slice was fixed with 4% PFA in PBS for 15 min at room temperate, followed by air dry and washing with PBS for 2 times, 5 min each time. Permeation of tumor slice was conducted in 0.1% Triton X-100 in PBS (PBST) for 1 hour, followed by 3% BSA in PBST blocking for 1 hour at room temperate. Primary antibodies (1:100 dilution in 3% BSA in PBST) were later added and incubated with the tumor slice overnight at 4°C. After 3 times PBS washing, secondary antibodies conjugated with fluorescent dyes (1:100 dilution in 3% BSA in PBST) were added and incubated with the tumor slice for 2 hours at room temperature. After 3 times PBS washing, the tumor slice was mounted and proceeded to Airyscan imaging. Details of the reagents used are listed in Key Resources Table.

#### Whole lipid and protein Expansion microscopy

0.0125x10^6^ cells were seeded and cultured overnight at a plasma-cleaned cover slip attached with a custom PDMS chamber (1 mm thickness and 6.5 mm diameter culture area, made with SYLGARD™ 184 Silicone Elastomer Kit) as previously described ^18^. To label the whole lipids (Figure 1D and 3A), cells were fixed with 37 °C pre-warmed 4% PFA and 0.1% Glutaraldehyde in PBS for 10 min, followed by washing with PBS twice and incubation with 5 µM mCling in PBS overnight at room temperature. After mCling staining, the cells were fixed again with 37 °C pre-warmed 4% PFA and 0.1% glutaraldehyde in PBS for 10 min then proceeded to standard immunostaining steps if required (Figure 3H and I). This included permeation with PBST, blocking with 3% BSA in PBST, and primary antibody incubation overnight at 4 °C. Secondary antibodies conjugated with Alexa Fluor 488 or Alexa Fluor 568, or secondary antibodies conjugated with LR-ExM linkers were used (1:50 dilution in 3% BSA in PBST) to fulfill multi-color ExM imaging and incubated with cells for 1 hour. Permeation step was still required to ensure isotropic expansion of ExM samples even though immunostaining was not applied. We found that glutaraldehyde can block the epitopes of proteins at nuclear lamina and nucleolus, so the fixation method for Figure 3B to G and Figure 4 A and B was 4% PFA in PBS for 10 min and mCling staining was not applied. To visualize F-actin, cells were stained with 1.65 µM phalloidin-fluorescein in 3% BSA in PBST for 1 h before primary antibodies incubation. To retain the phalloidin-fluorescein signal during expansion microscopy, cells were incubated with anti-fluorescein antibody conjugated with Alexa 488 dye (1:100 dilution in 3% BSA in PBST with other primary antibodies) overnight at 4 °C (Figure S7).

To conduct the expansion microscopy, the mCling-labeled and immunostained cells were further incubated with 0.04% (w/v) glycidyl methacrylate (GMA) in 100 mM sodium bicarbonate for 3 hours at room temperature, followed by washing with PBS for 3 times. GMA-treated cells were incubated with pre-chilled monomer solution (8.6 g sodium acrylate, 2.5 g acrylamide, 0.15 g N,N’-methylenebisacrylamide, 11.7 g sodium chloride in 100 ml PBS buffer) on ice for 5 min and later with gelation solution (mixture of monomer solution, 10% (w/v) N,N,N′,N′ Tetramethylethylenediamine stock solution, 10% (w/v) ammonium persulfate stock solution and water at 47:1:1:1 volume ratio) on ice for another 5 min. A cover slip was applied onto the top of the PDMS chamber to seal the cell-gelation solution to avoid oxygen interruption of the gelation procedure. The cells in gelation solution were later transferred to a 37 °C humidity chamber to initiate the gelation. After 1.5-hour gelation, the gelled cells were immersed in heat denaturation buffer (200 mM sodium dodecyl sulfate, 200 mM NaCl, and 50 mM Tris pH 6.8) for 1.5 hours at 78°C and washed with excess of PBS for 2 times, each time 30 min.

If secondary antibodies conjugated with LR-ExM linkers were used (Figure 3B, C, E and F), the gelled cells were first immersed in staining buffer (10 mM HEPES, 150 mM NaCl, pH 7.5) with anti-digoxigenin dye or streptavidin dye overnight, then proceeded to total protein labeling. To label the total proteins of cells (Figure 1D, 3A-C, E, F, and H-J), the gelled cells were shake and incubated with 20 µg/mL N-hydroxysuccinimide ester dye in PBS overnight. Finally, the gelled cells were immersed in great amount of DNase/RNase-free water and fully expanded at ∼3.8-4.1 times. The gelled cells were transferred to a poly-lysine-coated glass bottom dish prior to Airyscan imaging. Details of the reagents used are listed in Key Resources Table.

#### Nanopillar Fabrication

The fabrication of vertically aligned nanopillar arrays involved several steps. Initially, the silicon dioxide substrate was meticulously cleaned using water, acetone (Thermo Scientific, cat#268310025), and isopropyl alcohol (Thermo Scientific, cat#268310025)). A spin-coating process applied layers of poly(methyl methacrylate) (PMMA) (Allresist, cat#AR-P 672.045) and conductive polymer (Allresist, cat#AR-PC 5090.02) to the substrate surface. Using electron beam lithography (EBL) with a FEI Helios Nanolab 650, specific patterns were defined on the chip surface to determine the nanopillar’ size and pitch. The exposed PMMA was then dissolved in the developer (Allresist, cat#AR 600-56), followed by the deposition of 50 nm chromium (Cr) (LEE & LIM INTERNATIONAL) through thermal evaporation (UNIVEX 250 Benchtop). The redundant Cr was removed by lift-off using acetone, leaving the Cr mask with nanopillar patterns. Subsequently, reactive ion etching (RIE) (Oxford Plasmalab 80) utilizing CF4 and CHF3 gas was employed to achieve nanopillar’s height of 1.5 μm. Briefly, the CF4 and CHF3 gas was ionized by an electric field and formed plasma. The plasma was accelerated by a DC bias voltage and bombarded perpendicularly to the quartz substrate. Thus, the anisotropic reaction between the plasma and the glass ensured that the nanopillar had vertical sidewall and high aspect-ratio ^73^. The height of the nanopillar could be controlled by the electric field power, the gas pressure, and the reaction time.

Scanning electron microscopy (FEI Helios NanoLab 650) was utilized for characterization after another Cr coating. Finally, the Cr coating was washed with the chromium etchant solution (Sigma, cat#651826-500ML) and water, followed by nitrogen drying.

#### Cell culture on nanopillars

The cover glass with nanopillar arrays was first immersed in 98% sulfuric acid (Fisher Scientific, cat# 258105) overnight, followed by washing with excess amount of deionized water and air-drying. A PDMS chamber with 1 mm thickness and a 6.5 mm diameter hole was firmly attached to the nanopillar glass, carefully leaving the nanopillar area as culture area without contact of PDMS. The PDMS-chambered nanopillar glass was later cleaned in Harrick Plasma Basic Plasma Cleaner for 5 min with oxygen supply. Plasma-cleaned nanopillar glass was stored in the well of a sealed 12-well tissue culture plate until usage. 0.0125x10^6^ cells in a droplet of growth medium (∼50 µL) were seeded at the culture area of the nanopillar glass. Gently filled the well containing the nanopillar glass with 1 mL growth medium after 4 hours of cell seeding or until the cells attached to the nanopillar arrays. The cells attached to the nanopillar arrays were cultured overnight at 37 °C with 5% CO2 supply and later proceeded to RNA labeling and immunostaining.

#### Plasmids construction

GFP-HP1alpha was from Addgene (cat#17652). pSnap-LMNA was constructed via cloning LMNA (NM_170707.4) into pSNAPf-C1 vector with snap tag at the N terminus. pClip-LMNB1 was constructed via cloning LMNB1 (NM_005573.4) into pCLIP-tag (m) vector with clip tag at the N terminus. Plasmids for LaminA/C and LaminB1 knockout were ordered from VectorBuilder. The sgRNA #9142 sequenced TCGGGTCTCATGACGGCGCT and sgRNA#9143 sequenced GCGCCGTCATGAGACCCGAC were cloned into Cas9-puro-mCherry vector respectively for human LMNA knockout. The sgRNA #276 sequenced GTCGAGCGCGCGTCGCGCGT and sgRNA #277 sequenced GCGACGCGCGCTCGACGACA were cloned into Cas9-puro-mCherry vector respectively for human LMNB1 knockout. All plasmids used were full-plasmid sequenced and confirmed without errors.

#### Plasmids transfection

Before the plasmids transfection, cells were seeded at the density of 0.025x10^6^/well, cultured in 8 well glass bottom slide (ibidi, cat#80827) overnight, and starved with Opti-MEM medium (Invitrogen, cat# 31985062) for 30 min. For transfection of cells in one well of the 8 well glass bottom slide, 0.25 μg of plasmids were mixed with 0.5 μL of p3000 reagent in 12.5 μL Opti-MEM medium and added to 12.5 μL Opti-MEM medium with 0.75 μL of lipofectamine 3000 reagent (Invitrogen, cat# L3000001). The mixture was reacted for 15 min before addition to the cells. The cells were cultured with the plasmids mixture in Opti-MEM medium for 4 hours and later in growth medium without the plasmids mixture overnight. Cells transfected with the plasmids were proceeded to RNA labeling and immunostaining at the next day. Details of the plasmids used are listed in Key Resources Table.

#### Imaging

The imaging in this study was all performed on Airyscan confocal microscope (ZEISS LSM 980 with Airyscan 2) with a 63x water-immersion objective (Zeiss LD C-Apochromat 63x/1.2 W Corr M27) with effective lateral resolution at 138 nm (measured by TetraSpeck™ Microspheres, 0.1 µm, fluorescent blue/green/orange/dark red). Airyscan SR-4Y and best signal mode with 0.2 AU pinhole and 1.25 AU total detection area were used for the 3D imaging of all the samples. After combination with expansion microscopy, the actual lateral resolution was enhanced to ∼35 nm. For live cell imaging, a stage-top incubator system (ibidi, cat#12720) was installed and applied. 0.22x10^6^ cells were seeded at 35 mm glass bottom dish (Mattek, P35G-1.5-14-C) and cultured overnight before the day of imaging. Airyscan SR-4Y with imaging acquisition speed at ∼800 milliseconds was applied.

#### Western blotting

To compare the ribosome biogenesis level in MCF-10A and MDA-MB-231 cells, two groups of MCF-10A and MDA-MB-231 cells were seeded (0.22x10^6^/well) and cultured overnight in the wells of 6-well plate. A group of cells were trypsinized and dissociated for cell counting at the day of collection. The other group of cells were placed on ice and lysed with 1x laemmli sample buffer (beta mercaptoethanol added) (Bio-Rad, cat# 1610747) for western blotting. The cell lysates were sonicated and centrifuged before SDS-PAGE gel loading. Equal amount of cell lysates with same number of cells were loaded to the TGX stain-free protein gels (Bio-Rad, cat# 4568124). Stain-free total protein imaging was performed on ChemiDoc MP imaging system (Bio-Rad, cat#12003154). After transfer, the PVDF membrane was blocked with 3% BSA in TBST (20 mM Tris-HCl, pH 7.5, 150 mM NaCl, 0.05% Tween-20) for 30 min and incubated with primary antibodies (1:1000 dilution in 3% BSA in TBST) overnight at 4 °C. After the overnight incubation, the membrane was washed with TBST for 3 times (10 min/time) and incubated with HRP-conjugated secondary antibody (1:10000 dilution in TBST) for 1 hour at room temperature. After the secondary antibody incubation, the membrane was washed with TBST for 3 times (10 min/time) and proceeded to imaging with Clarity Max™ Western ECL Substrate (Bio-Rad, cat# 1705062) at ChemiDoc MP imaging system. Details of the antibodies used are listed in Key Resources Table.

#### Modeling of ribosome biogenesis and export

To model the pre-ribosome biogenesis and transport through the nucleus, we developed a partial-differential-equation (PDE) model of the concentration of pre-ribosomes in the nucleus. The model encodes diffusive motion and export out of the nuclear boundary. Diffusion is assumed to vary depending on heterochromatin. Export of pre-ribosomes out of the nucleus is modeled by a semi-permeable boundary, with an absorption parameter encoding NPC density and export delay. The output of the model is the overall flux out of the nucleus scaled by the Exportin1 concentration based on the assumption of a first-order binding. Exportin1 concentration, NPC density, heterochromatin thickness, nucleolar size, and nucleolar pre-ribosome density all may vary with pillar radius based on measured quantities. The model output is computed for various pillar radii, normalized by the R350 output flux so it may be compared directly with mature ribosome measurements. The model is solved numerically using MATLAB’s finite element PDE Toolbox, and the parameters are fitted using a mean-squared error against the mature ribosome data for different pillar radii. See **Supplementary Information** for further information.

### Quantification and statistical analysis

#### Images analysis

All images were processed and analyzed using ImageJ and Custom MATLAB code.

#### Quantification of nuclear invaginations

3D stacks of LaminB2 images were max intensity projected into 2D images. Threshold function with default setting in imageJ was first applied to every single nucleus in the 2D images to generate individual region of interest (ROI) that outlined the entire nucleus. A second threshold function with default setting was applied to the selected nuclear invagination area in the 2D images. The ratio of nuclear invagination area to the entire nucleus area was measured in every single nucleus. This ratio was used as measurement of nuclear invagination level.

#### Quantification of nucleolar EU intensity and RPA194 intensity

3D stacks of EU images were max intensity projected into 2D images. Threshold function with default setting in imageJ was applied to the selected nucleolar area in the 2D images and tracing tool was used to generate the outline of each nucleolus. The mean EU intensity and mean RPA194 intensity within every single nucleolus were measured. Same method was applied on measuring the RPL13 intensity inside single cell, eIF6, H3K9me3, Lamin intensity inside single nucleus.

#### Quantification of size of nuclear invaginations and heterochromatin thickness

3D Airyscan-Expansion microscopy images of whole nucleus were taken as shown in Movie S1. The scale of the 3D image was set to its pre-expansion unit based on the length expansion factor measured and the actual size of the image taken. Since the diameter of single nuclear invagination was various, the smallest diameter of each nuclear invagination at the NE-nucleolus contact was measured via the straight-line tool and measure function in imageJ (Figure 1I). Similarly, smallest heterochromatin thickness was measured at each NE-nucleolus contact (Figure 4Aiii and Figure 4Biii, line with both flat ends).

#### Quantification of NPC numbers and RPA194 cluster

The effective lateral resolution of our Airyscan-Expansion microscopy was around 35 nm, which is smaller than half diameter of single NPC (∼120 nm). Therefore, our Airyscan-Expansion microscopy can resolve single NPC. To count the number of the NPC at the NE-nucleolus contact, we used Nup153, which is located at the basket of NPC with diameter around 50 nm. That means each dot in the Nup153 images represent one NPC (Figure 4Aiii and Figure 4Biii). The length of the NE-nucleolus contact was measured via freehand selection tool of imageJ. The number of NPC on the NE-nucleolus contact was counted and the density of NPC was subsequently calculated by number of NPC versus length of NE-nucleolus contact (Figure 4D).

Our Airyscan-Expansion microscopy can also resolve single FC region inside the nucleolus, as shown in Figure 3A and Movie S1. RPA194 was shown as a cluster inside the FC region of nucleolus (Figure 3B and 3C). To count the number of RPA194 cluster, 3D stacks of RPA194 images and nucleolus images (NHS staining was used to image the nucleolus) were max intensity projected into 2D images. The area of each nucleolus was measured via freehand selection tool of imageJ. And the number of RPA194 cluster was counted. The density of RPA194 cluster was subsequently calculated by number of RPA194 cluster versus area of nucleolus (Figure 3D).

#### Quantification of lipid and protein distribution at the nanopillars

Image processing and analysis were performed using custom-written MATLAB (Mathworks) code adapted from previous work ^22^. Briefly, artificial nuclear tunnel generated by single nanopillar was located using laminA/C, laminB1 or mCling channel and individual tunnel image was cropped by a square mask (71×71 pixels) centered at the nuclear tunnel. Background of each individual tunnel image was corrected by subtracting the mean intensity of four ROIs (10×10 pixels) at the corners of the image. Background-corrected individual tunnel images with same nanopillar radius were then averaged across different arrays and experimental repeats to generate averaged images (as shown in Figure 4E). To quantify the lipid and protein density at artificial nuclear tunnels, each tunnel image was segmented into two ROIs: tunnel center (a small ROI that covered the nanopillar area without lipid and protein signal) and tunnel edge (a large ROI including the tunnel center and the lipid and protein signal around the nanopillar). The sizes of the tunnel center and tunnel edge were adjusted based on the dimension of the nanopillar and signals around the nanopillar. Lipid and protein density at the tunnel were subsequently calculated by (integrated intensity at the tunnel edge – integrated intensity at the tunnel center)/perimeter of nanopillar.

## Statistical analysis

P values were determined with Student’s t tests and all graphs were generated using Prism 10 (GraphPad software).

## Acknowledgments

We acknowledge Dr. Wenqi Wang at UC Irvine, Dr. Pablo Lara-Gonzalez, Dr. Fangyuan Ding at UC Irvine, and members of the Shi lab for the helpful discussions and advice. We thank Dr. Jie Chen, Dr. Zhipeng Dai, and Adrian Chao for the technical support on the expansion microscopy imaging. We thank Dr. Alana Welm at University of Utah for providing HCI-002 PDX and Dr. Jordan Woytash for sharing frozen HCI-002 PDX tumors. Y.Z. is supported by an NSF-Simons grant (DMS1763272) and K99/R00 NIH Pathway to Independence Award (R00GM126136). O.V.R. is supported by 25IB-0059 California Breast Cancer Research Program (CBCRP). C.E.M. is supported by NSF Division of Mathematical Sciences Award (2339241). X.G. and W.Z. are supported by fundings from the Singapore Ministry of Education (RG145/18, NGF-2021-10-026 and MOE-MOET32020-0001), the Singapore Ministry of Health (MOH-001192-01), Human Frontier Scientific Program (RGY0088/2021), and Nanyang Technological University (Start-Up Grant). X.S. is supported by the NIH Director’s New Innovator Award (DP2GM150017) and NSF Faculty Early Career Development Program (CAREER) Award (2341058). The project is also supported by the Chan Zuckerberg Initiative (CZI) Visual Proteomics Imaging Award and the Advancing Imaging Through Collaborative Projects Award. The project was initiated with a pilot grant from the UCI Center of Cancer Systems Biology Grant (U54-CA217378).

## Author contributions

Conceptualization: YZ, XS

Methodology: YZ, XS, WZ, XG, OR, CM

Investigation: YZ, XS, CM

Visualization: YZ, XS

Funding acquisition: XS, WZ, OR

Project administration: YZ, XS

Supervision: XS

Writing – original draft: YZ, XS, CM

Writing – review & editing: YZ, XS, WZ, XG, OR, CM

## Declaration of generative AI and AI-assisted technologies in the writing process

During the preparation of this work, the authors used ChatGPT for language and grammar checks. After using this tool, the authors reviewed and edited the content as needed and took full responsibility for the publication’s content.

