## Supplemental Information for "Coaching ribosome biogenesis from the nuclear periphery"

**Affiliations:**
[1]Department of Developmental and Cell Biology, University of California, Irvine; Irvine, CA 92697, United States.
[2]School of Chemistry, Chemical Engineering and Biotechnology, Nanyang Technological University; Singapore 637459, Singapore.

[3]Department of Molecular Biology and Biochemistry, University of California, Irvine; Irvine, CA 92697, United States.

[4]Department of Mathematics, University of California, Irvine; Irvine, CA 92697, United States.

[5]Department of Chemistry, University of California, Irvine; Irvine, CA 92697, United States.

[6]Department of Biomedical Engineering, University of California, Irvine; Irvine, CA 92697, United States.

[7]Lead contact

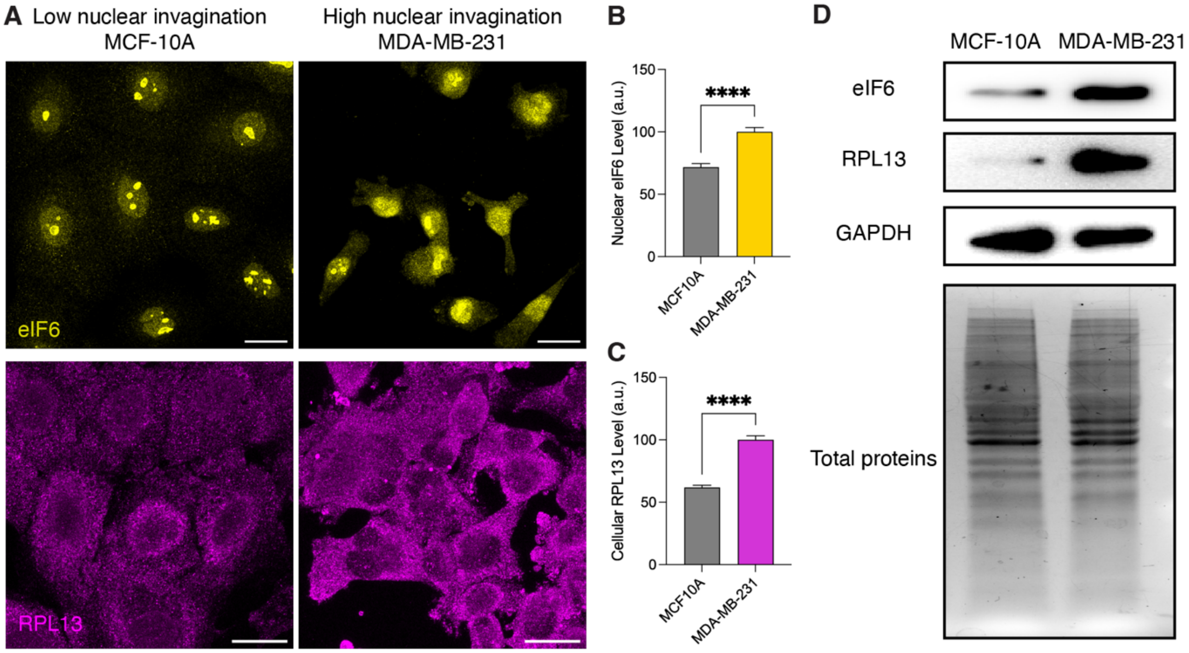


***Figure S1. Nuclear Invagination is Associated with Ribosome Biogenesis.***

*(A) Airyscan images of MCF-10A and MDA-MB-231 cells immunostained with anti-eIF6 (yellow) and anti-RPL13 (magenta) antibodies. Scale bars: 20 µm.*

*(B) Analysis of eIF6 intensity within the nuclei of MCF-10A and MDA-MB-231 cells. Each bar represents mean ± standard error of more than 20 cells. **** indicates p<0.0001 by unpaired t test.*

*(C) Analysis of RPL13 intensity of MCF-10A and MDA-MB-231 cells. Each bar represents mean ± standard error of more than 35 cells. **** indicates p<0.0001 by unpaired t test.*

*(D) Western blot images of MCF-10A and MDA-MB-231 total protein extracts and membrane immunostained with anti-eIF6, anti-RPL13 and anti-GAPDH antibodies.*

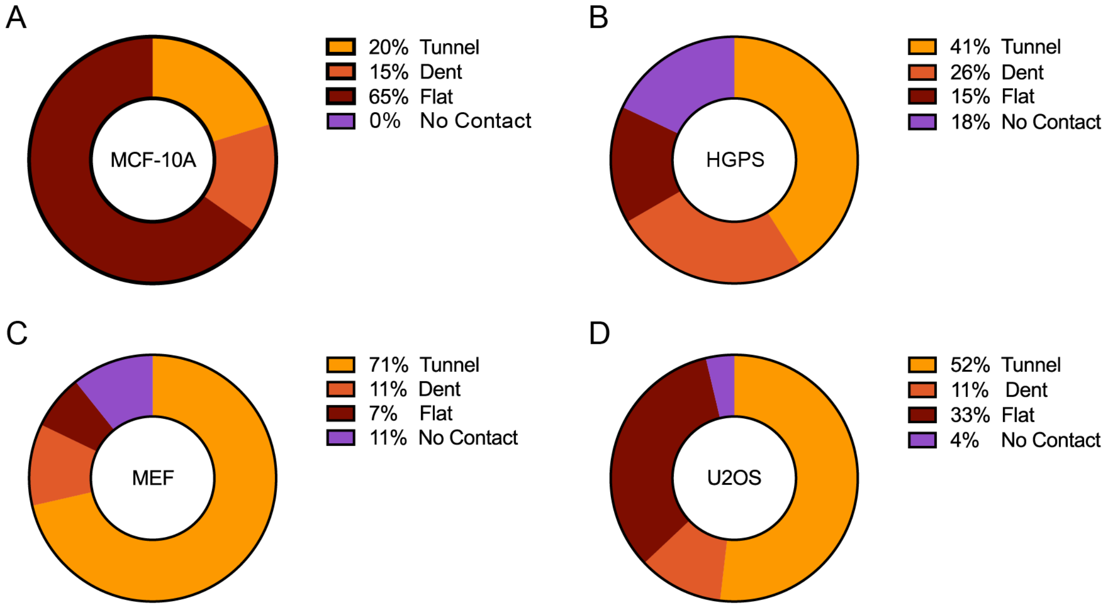


***Figure S2. Probability of Different types of NE-Nucleolus Contacts in Cell Lines.*** *69, 39, 56, 54 nucleoli were analyzed in MCF-10A cells (A), progeria HGPS primary cells (B), MEF cells (C) and U2OS cells (D), respectively.*

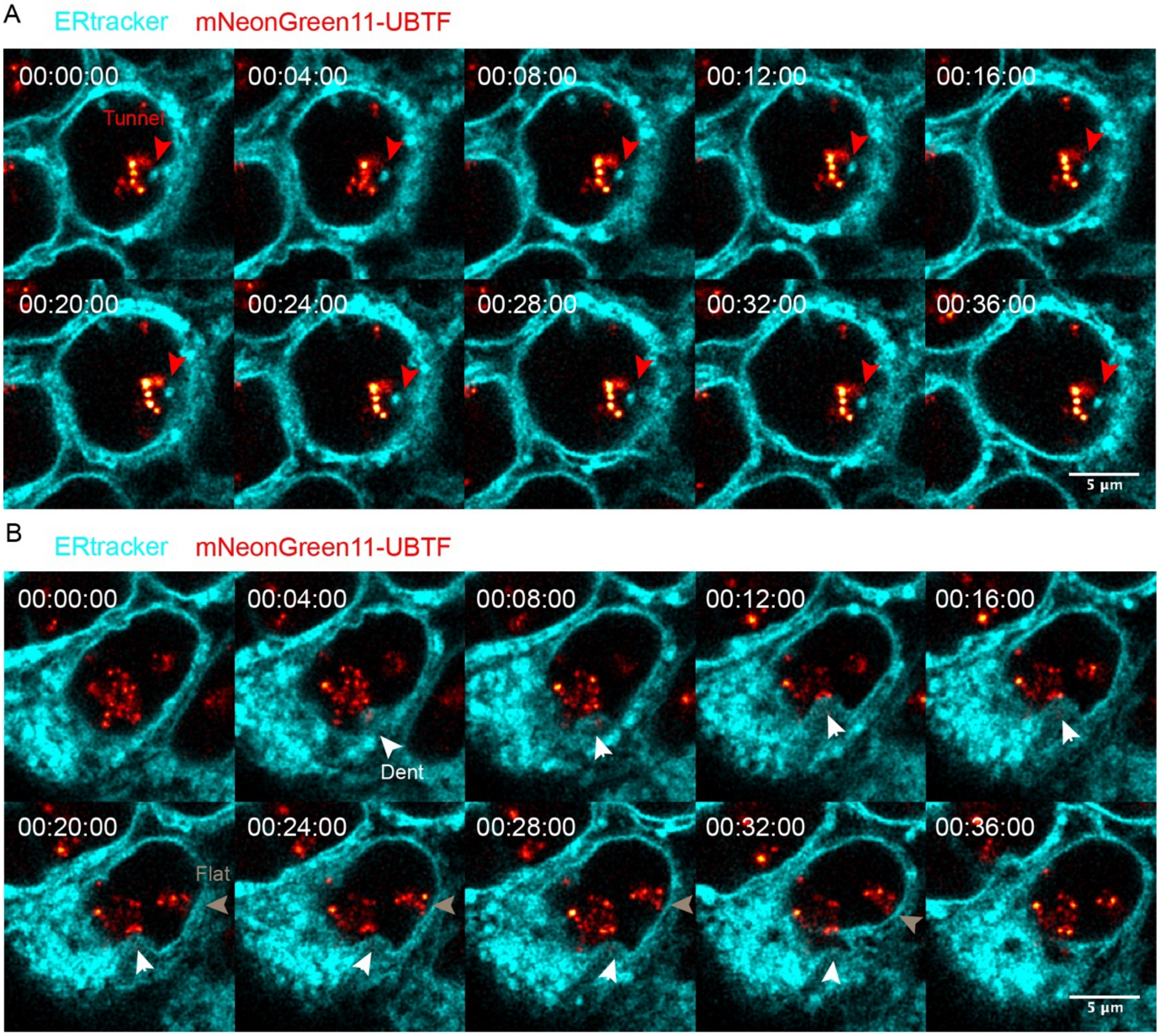


***Figure S3. Dynamics of NE-Nucleolus Contacts.***

*Airyscan time-lapse images of HEK293T cells CRISPR knocked in split mNeongreen-UBTF (red hot) and live-stained with ERtracker red (cyan). (A) Red arrows point to a nuclear tunnel-nucleolus contact site. (B) White arrows point to a nuclear dent-nucleolus contact site. Grey arrows point to a flat NE-nucleolus contact site. Scale bars: 5 µm.*

### *Cell Cultured on Nanopillar Patterns*

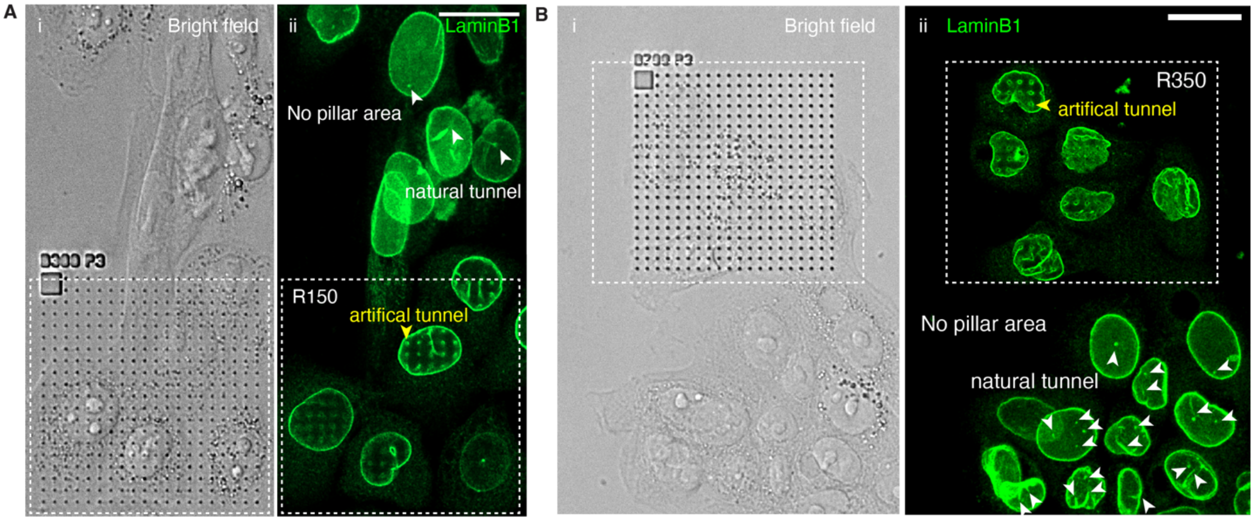


***Figure S4. Nanopillars induce nuclear invaginations of desired curvatures.***

*Brightfield and fluoresce two-color images of MCF-10A cells cultured on substrates with a nanopillar array and a flat surface region. The dashed line boxes outline the nanopillar areas. All cells were immunostained with anti-LaminB1 antibodies (green). All images were taken on an Airyscan microscope. (A) Images of cells cultured on a substrate containing nanopillars with a 150 nm radius. i: Brightfield channel. ii: LaminB1 fluorescence channel. (B) Images of cells cultured on a substrate containing nanopillars with a 350 nm radius. i: Brightfield channel. ii: LaminB1 fluorescence channel. White arrowheads point to the natural tunnels and yellow arrowheads point to artificial tunnels generated by the nanopillars. Scale bars: 20 µm.*

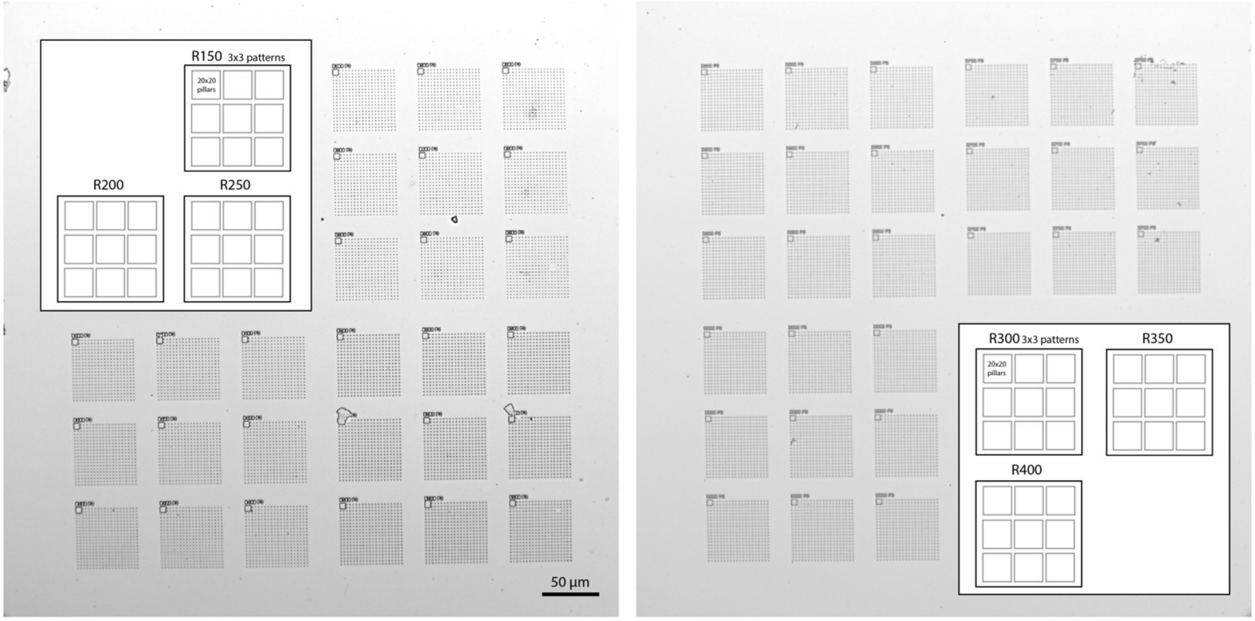

***Figure S5. Full View of Nanopillar Patterns.*** *Brightfield images of nanopillar patterns with radii from 150 nm (R150) to 400 nm (R400). Scale bar: 50 µm. Inserts: Blueprints of the nanopillar patterns. On one quartz, there are 3 groups of nanopillars with different radii. Each group consists of 9 patterns. Each pattern consists of 20x20 nanopillars.*

#### ***Mitochondria Interactions at Nuclear Tunnel***

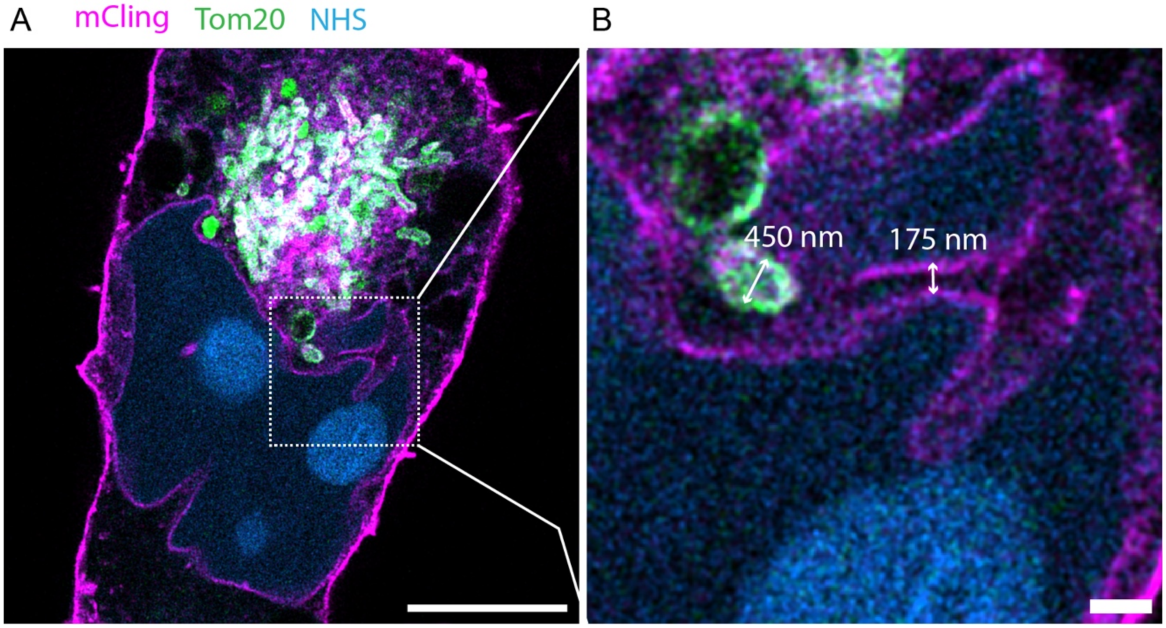


***Figure S6. Mitochondria in the nuclear tunnel.*** *(A) Airyscan expansion microscopy image of UCI082014 cell stained with anti-Tom20 (green) antibodies, NHS ester (blue), and mCling (magenta). Scale bar: 5 µm in pre-expansion unit. (B) Magnified view of the boxed area of image (A). Scale bar: 500 nm in pre-expansion unit. Length expansion factor: 3.7.*

#### ***F-actin Puncta inside the Nuclear Tunnel***

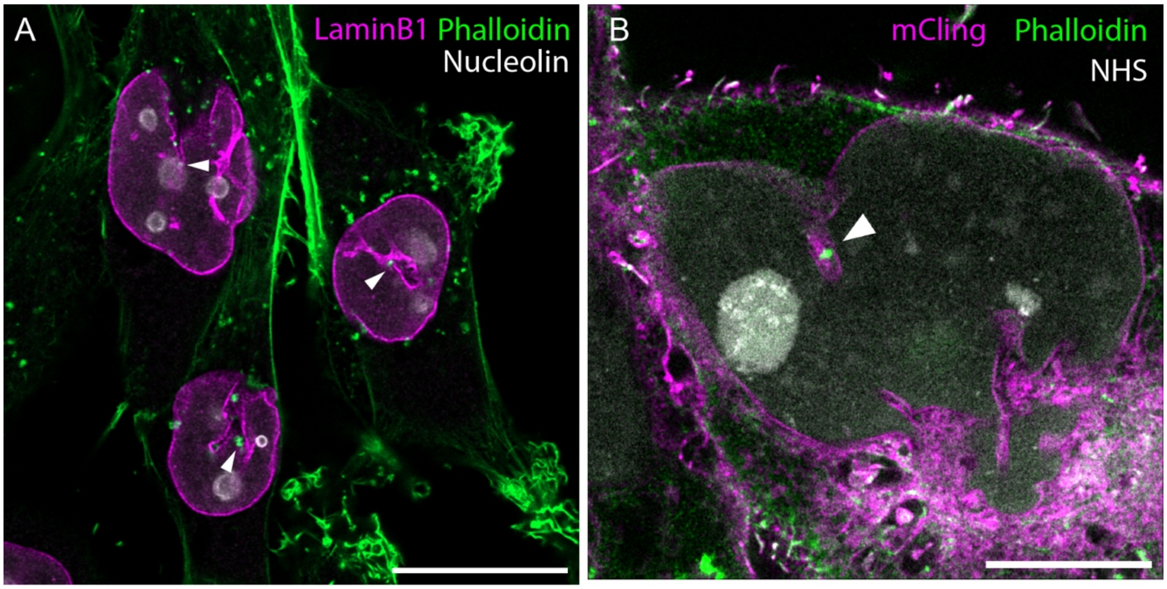


***Figure S7. F-actin puncta inside nuclear tunnels.*** *(A) Airyscan image of UCI082014 cells stained with phalloidin-FITC (green), anti-LaminB1 antibodies (magenta), and anti-nucleolin antibodies (grey). Scale bar: 20 µm. (B) Airyscan expansion microscopy image of UCI082014 cells stained with phalloidin-FITC and anti-fluorescein antibody (green), NHS ester (grey), and mCling (magenta). Scale bar: 5 µm in pre-expansion unit. White arrows point to F-actin puncta. Length expansion factor: 3.8.*

***mTOR Signaling and Nuclear Invaginations***

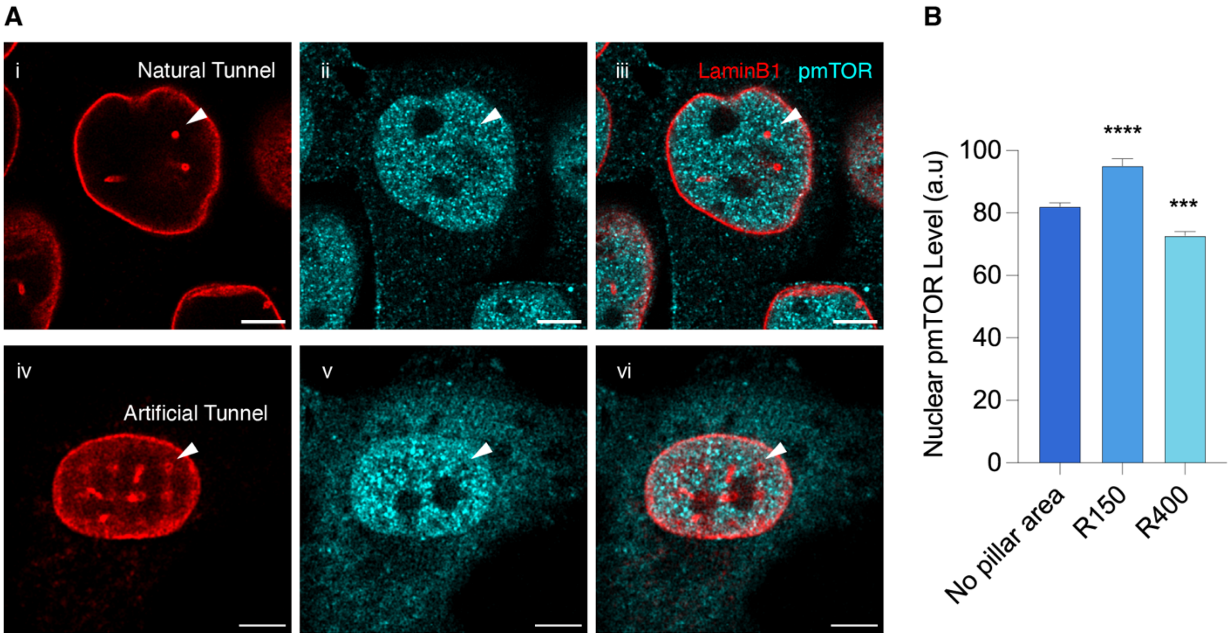


***Figure S8. Nuclear phosphorylated-mTOR is Curvature-Dependent.***

*(A) (i-iii): Airyscan images of UCI082014 cells immunostained with anti-LaminB1 (red) and anti-phosphorylated-mTOR (cyan) antibodies. White arrowhead indicates the natural nuclear tunnel. The black holes inside the nucleus at the pmTOR channel shape nucleoli. (iv-vi): Airyscan images of MCF-10A cells seeded on the pillars with 300 nm radius immunostained with anti-LaminB1 (red) and anti-phosphorylated-mTOR (cyan) antibodies. Arrowhead indicates artificial tunnels generated by the nanopillars. The black holes inside the nucleus at the pmTOR channel shape nucleoli. Scale bar: 5 µm.*

*(B) Barchart of phosphorylated-mTOR intensity in nuclei of MCF-10A cells cultured on flat surfaces and nanopillars with radii of 150 nm (R150) and 400 nm (R400). Each bar represents mean ± standard error of more than 40 nuclei. **** indicates p<0.0001 and *** indicates p<0.001 by unpaired t test.*

***Diffusion Model Description***

**Full two-dimensional model**

For a prescribed nuclear and nucleolar geometry (see section **Geometry Generation**), we assume a diffusion model for the motion of the pre-ribosomes between the nucleolus and export out of the nucleus through NPCs. This is based on single-molecule measurements that suggest this motion is indeed diffusive, or at least well-approximated [1,2].

For parameters that may depend on the particular geometry, we denote these as functions of $R$, where we consider $R$ to index the pillar radii $R = \{150, 250, 300, 350, 400\}$ in nm.

The concentration of pre-ribosomes $u(\vec{x}, t)$ is assumed to follow Fickian diffusion

$$\partial_t u = \nabla \cdot D(\vec{x}) \nabla u, \qquad (1)$$

where we assume that creation and export are in equilibrium, so we consider steady-state (setting $\partial_t u = 0$), yielding

$$0 = \nabla \cdot D(\vec{x}) \nabla u. \qquad (2)$$

The diffusion of pre-ribosomes is assumed to differ in heterochromatin regions and other nuclear space

$$D(\vec{x}) = \begin{cases} D_{\text{slow}} & \vec{x} \in \text{heterochromatin} \\ D_{\text{fast}} & \text{otherwise.} \end{cases} \qquad (3)$$

We also assume that pre-ribosomes are released in a non-depleting manner from the nucleolar surface with fixed concentration that may vary based on geometry,

$$u|_{\vec{x} \in \text{nucleolus boundary}} = p(R). \qquad (4)$$

Without direct knowledge of the explicit location of the NPCs, we instead follow and model export through the NPCs using a surface reactivity (Robin condition) term [3,4]

$$\vec{n} \cdot (D(x)\nabla u) + k(\vec{x}, R)u|_{\vec{x} \in \text{nucleus boundary}} = 0. \qquad (5)$$

Here, $\vec{n}$ is the outward normal vector and $k(\vec{x}, R)$ describes the surface reactivity, which may vary between the pillar geometries. The spatial dependence arises from the observation that NPC density varies between the pillar and non-pillar flat portions of the nuclear surface, modeled as

$$k(\vec{x}, R) = \begin{cases} k_{\text{pillar}}(R) & \vec{x} \in \text{pillar surface} \\ k_{\text{flat}} & \text{nuclear surface otherwise.} \end{cases} \qquad (6)$$

Interpreting this boundary condition: as $k$ becomes large, pre-ribosomes exit through NPCs more quickly. Alternatively, whenever a pre-ribosome diffuses into the nuclear surface, $k$ captures the probability of how likely it is to be exported [5]. Ultimately, this term serves to approximate (1) the known delay in export through an NPC [6] and (2) NPC distribution on the nuclear surface.

These PDEs are solved using MATLAB's Finite Element PDE Toolbox. From the solution, the flux through the boundary is computed by

$$\text{boundary flux} = -\int_{\vec{x} \in \text{nucleus boundary}} \vec{n} \cdot \nabla D(\vec{x}) u(\vec{x})\, \mathrm{d}\vec{x}, = \int_{\vec{x}, \in \text{nucleus boundary}} k(\vec{x})\, u(\vec{x})\, \mathrm{d}\vec{x} \qquad (7)$$

At the stage of this surface integral, we reiterate acknowledgement that this two-dimensional approximation does not faithfully reflect the full three-dimensional nature of the system and presumably has different scaling properties. However, we seek to understand relative trends in the flux, the exploration of which is possible because of this simpler setup.

We next note that exportin is required for a pre-ribosome to be exported through an NPC. In lieu of resolving bound and anchored exportin, we assume the binding process between pre-ribosomes and exportin is a fast, first-order reaction. That is, pre-ribosomes bind with exportin to form a complex $P + X \leftrightarrow C$ with a fixed pool $E$ of exportin where $X + C = E$, then in equilibrium $C \propto E$. Noting that the complex is exported, we use this argument to scale the flux by the exportin concentration. Due to the observation that exportin densities vary between the pillar and non-pillar regions (flat) on the nuclear surface, we also incorporate this dependence.

The resulting final expression that is compared with mature ribosome measurements in the main text is

$$J = \text{flux}_{\text{pillar}} E_{pillar}(\mathrm{R}) + \text{flux}_{\text{flat}} E_{flat}$$
$$= E_{pillar}(R) \int_{\vec{x} \in \text{pillar boundary}} k_{\text{pillar}}(R)\, u(\vec{x})\, \mathrm{d}\vec{x}$$
$$+ \; E_{flat} \int_{\vec{x} \in \text{flat boundary}} k_{\text{flat}}\, u(\vec{x})\, \mathrm{d}\vec{x}\,. \qquad (8)$$

This expression describes the "Full Model" predictions in the main text.

**Approximate Model**

To understand the behavior of the model more intuitively, we note that since we are only interested in the relative flux change for different conditions, we can instead consider the pointwise flux at the boundary rather than the integrated total flux. Therefore, we consider the 1-dimensional approximation where $x = 0$ is the nuclear surface (with NPCs) and $x = L$ is the nucleolar surface,

$$0 = \partial_x D_{\text{slow}}\, \partial_x u, \quad -D_{\text{slow}}\, \partial_x u(0) + ku(0) = 0, \quad u(L) = p. \qquad (9)$$

The distance $L$ from the nucleolus to the pillar is assumed to be the measured heterochromatin thickness. Moreover, we assume that diffusion at the nuclear surface is through heterochromatin, so $D = D_{\text{slow}}$. This can be solved analytically using straightforward integration,

$$u(x) = p\frac{D_{\text{slow}} + kx}{D_{\text{slow}} + kL}. \qquad (10)$$

Finally, we make the same assumption about exportin scaling, yielding the analytical expression for the flux

$$J(r) = \text{flux} = E(R)ku(0) = E\frac{D_{\text{slow}}k(R)}{D_{\text{slow}} + kL(R)}p(R). \qquad (11)$$

This expression describes the "Simplified Model" predictions in the main text.

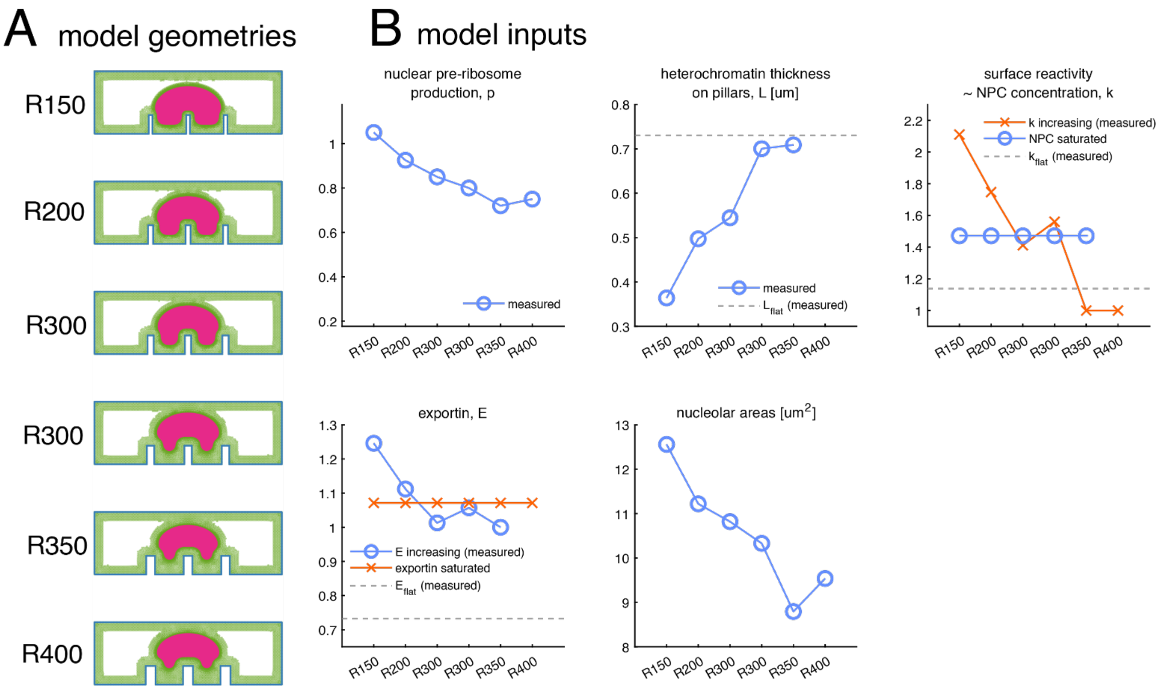


***Figure S8: Inputs to the model. A:*** *Model geometries used to solve the full 2D model. Green regions indicate (slow diffusion) chromatin regions. Pink shows the nucleolus. See "Geometry Generation" for details on generation.* ***B:*** *Pillar roundness dependency for each parameter. In the main text, k saturating (no R dependence) and E decreasing are used due to being the best fit with reasonable parameters (see "Model Fitting"). Values for R400 are taken to be the same as R350 if unmeasured.*

### Model fitting and variations

Only relative changes in mature ribosome can be measured, so all flux values are measured relative to $J(R = 350)$, and therefore we set $J(R = 350) = 1$. Then, the remaining values, e.g., the relative flux in the 150 nm pillar configuration $J(R = 150)/J(R = 350)$ are compared with the mature ribosome measurements, and fit with respect to MSE (mean-squared error) $\text{MSE} = (1/5)\sum_{R=\{150,200,250,300,350\}}[\text{RPL13}(R) - J(R)/J(350)]^2$ and this quantity is minimized over the parameters $D_{\text{slow}}$ and $\kappa$. Notably, since this is steady-state, all rates can be rescaled to yield the same solution, so $D_{\text{fast}} = 10$ is fixed, based on the observation of a free-cytosol diffusion coefficient of approximately 10+ $\mu m/s^2$ in previous studies [2,7].

In the "Full Model" (and the simplified 1D version) we assume $p, k, E$ and the heterochromatin thickness all vary with the pillar radius $R$ and scale directly proportionally to the observed measurements, as shown in ***Figure S8B***. Specifically, only relative values are shown, so the surface reactivity is taken so that $k(R) = \kappa \cdot \text{Nup153}(R)$ , where $\text{Nup153}(R)$ are the NPC measurements and $\kappa$ determines the overall magnitude of the process. These are again relative to the measurement at 350 nm pillar radius, so $\text{Npc153}(R = 350) = 1$.

We also consider several model variants. Both exportin1 and Nup153 concentrations should likely appear as Michaelis-Menten-like scaling in the model. However, in lieu of introducing this complexity, we instead investigate the two effective regimes: a low-concentration linear dependence and a saturating non-dependence. In this latter saturating scenario, the NPC import and exportin terms become independent of $R$, so we set $k(R) = \kappa$ or $E(R) = E = 1$. Each of these variants is shown in ***Figure S8B.***

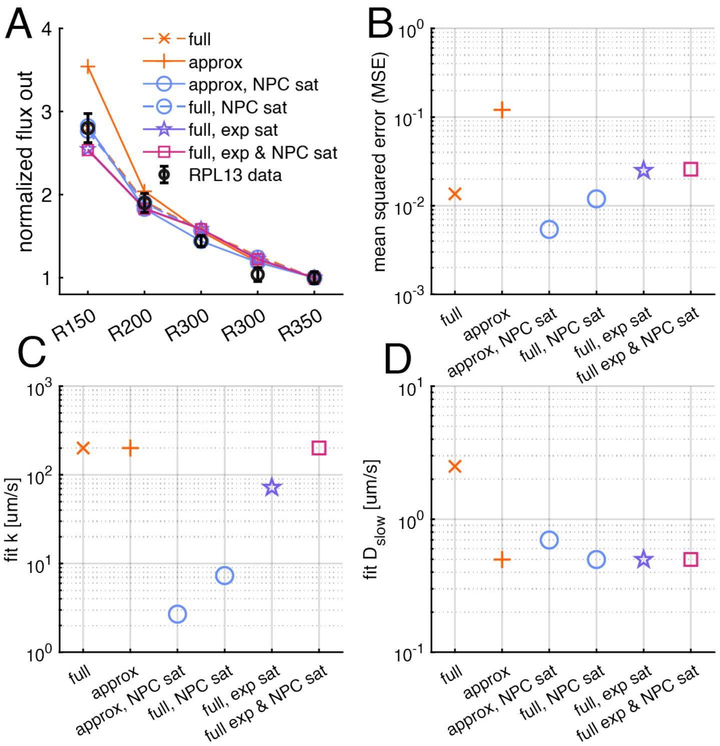


***Figure S9: A:*** *Fits for various competing models.* ***B:*** *Mean squared errors (MSE) for fits shown in the previous panel.* ***C:*** *Diffusivity fit for each scenario, and* ***D:*** *surface reactivity fit.*

The fitting procedure produces a notably good fit for all models, as shown in ***Figure S9A.*** However, we note that the scenarios (both full and approx.) with NPC saturation provide a discernibly better fit (***Figure S9B***). Moreover, these scenarios correspond to parameter regimes where both *D* and *k* have values in agreement with the literature (***Figure S9C and D*****).** From this, we conclude that export rate saturating in NPC density is the most plausible model, and the one used elsewhere unless noted otherwise.

***Role of Parameters***

As one last investigation of the model and corresponding fits, we take the model fits from the previous section and sweep over *D* and *k* to see how increases (or decreases) in these parameters shape the resulting flux. Specifically, we consider the NPC saturating scenario in the approximate model and take each parameter to be 100x smaller to 100x larger than its baseline (best fit) value, shown in ***Figure S10.***

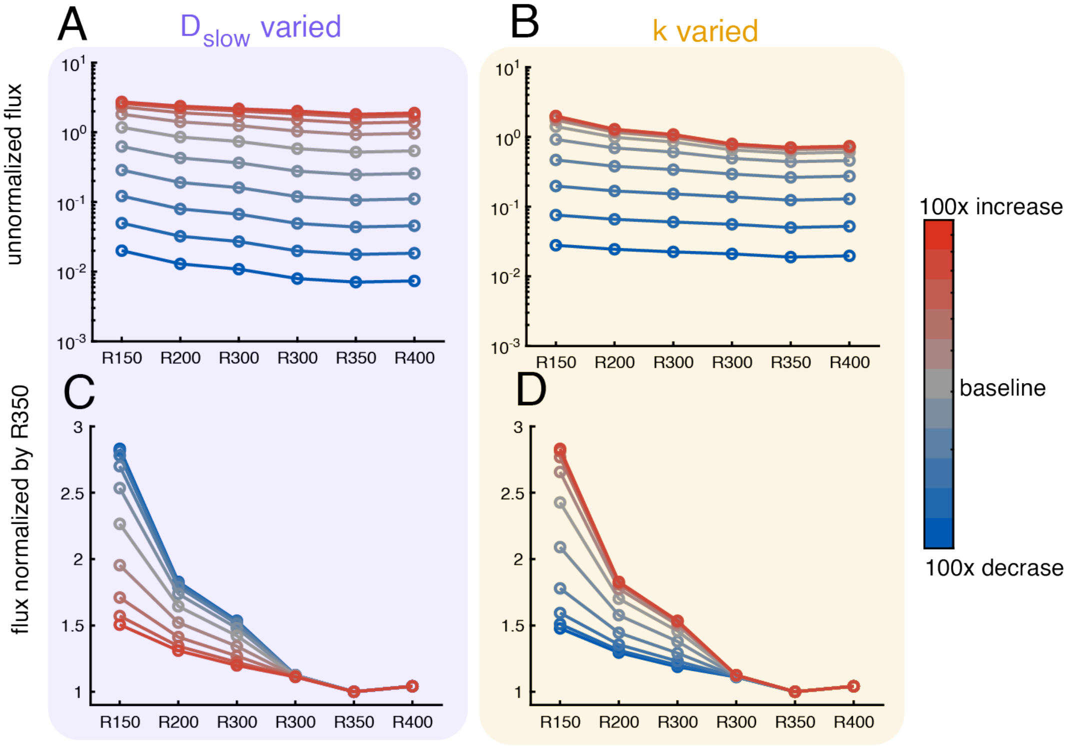


***Figure S10:*** *Role of D and k on output flux.* ***A:*** *Unnormalized (raw output of the model) flux from sweeping D in the approximate model. Larger D yields larger flux.* ***B:*** *Same plot and trend for sweeping k.* ***C:*** *Normalizing the curves in* ***A*** *by their value at R350, higher values of D flatten the flux curve.* ***D:*** *Normalizing by R350 reveals that larger values of k steepen the flux curve.*

In ***Figure S10A and B***, we see that increasing *D* or *k* yields the intuitive result that the overall flux increases when these parameters increase. However, the experimental data is normalized by the value at R350, and all measurements are relative to this. In ***Figure S10C and D*****,** the previous curves are normalized by their values at R350 and a surprising difference emerges. As *D* increases, the effect for different pillar radii gets washed out. In contrast, larger values of *k* yield larger differences in the flux for different pillar radii. This difference can be understood as follows: if diffusion through chromatin becomes fast, the varied thickness seen becomes increasingly negligible, and the curve flattens. However, for $k$ larger, meaning faster NPC export, the only limiting factor is diffusive transport time through the chromatin, so the varied thickness becomes the dominant difference. Altogether, this further emphasizes the main text point that export is a fine balance between these two competing timescales.

***Geometry Generation***

To generate the nucleolus geometries used in the diffusion model, we follow the methods of Ref. [8] to evolve a deformable solid experiencing a variety of biophysical forces. In contrast to that work, we emphasize that our forces are fictitious, and solely used to generate plausible geometries rather than trying to faithfully describe the complex physics of the nucleolus.

Each nuclear domain is first taken to be a $W$ by $H$ rectangle, approximating the side view of an approximately cylindrical nuclear domain. At the bottom of the rectangular domain, 3 rectangular pillars are added symmetrically around the center of the boundary, with varied pillar width, spacing 3 $\mu$m, and heights 1.5 $\mu$m to mirror setups.

The nucleus is initially configured at (fictitious) $t = 0$ to be a circle with radius $r$ specified so that the area matches experimentally observed values. The surface of the nucleolus is a parameterized curve $\vec{x}(s)$ with $0 \leq s \leq 1$. This curve is discretized into $M$ points, $\vec{x}_1, \dots, \vec{x}_M$ where we can associate $\vec{x}_{M+1} = \vec{x}_1$.

The points on the nucleolus surface then evolve via the sum of the forces exerted on them until apparent convergence $\| \partial_t \vec{x}_i \| < 10^{-6}$ or until $t = 1000$. The forces are in total,

$$\partial_t \vec{x}_i = \vec{F}_{\text{elast}} + \vec{F}_{\text{press}} + \vec{F}_{\text{down}} + \vec{F}_{\text{steric}}. \qquad (12)$$

**Elastic**: An elastic force on the surface of the nucleolus resists deformations caused by the other forces, and results in effective Hookean springs between the discretized points on the surface:

$$\vec{F}_{\text{elast}} = f_{\text{elast}}\, \partial_s (\| \partial_s \vec{x} \| - \ell)\vec{\tau}, \qquad (13)$$

where $\ell$ is the rest length of each connection determined by the initial configuration, $\vec{\tau} = \partial_s \vec{x} / \| \partial_s x \|$ is the tangent vector, and $f_{\text{elast}}$ controls the overall magnitude of this force. All $\partial_s$ derivatives are computed by a finite difference approximation $\partial_s \vec{x}_i \approx (\vec{x}_i - \vec{x}_i)/(M-1)$.

**Pressure**: To maintain the overall area of the nucleolus after deformations, a pressure force is exerted

$$\vec{F}_{\text{press}} = -f_{\text{press}} \ln(A/A_0)\vec{n}, \qquad (14)$$

where $A$ is the area enclosed by $\vec{x}$, $A_0$ is the initial area, and $\vec{n}$ is the outward normal vector.

**Downward force**: Based on the observation that the nucleolus deforms around the pillars and heterochromatin, a fictitious downward body force is exerted on the nucleolus so that it comes in contact with the pillars. This is taken to be

$$\vec{F}_{\text{down}} = -f_{\text{down}} \vec{y}, \qquad (15)$$

where $\vec{y} = [0,1]$, the vertical direction.

**Steric force**: Finally, the nucleus, pillar, and heterochromatin exert a steric force on the nucleolus itself, of the form

$$\vec{F}_{\text{ster}} = -f(\| x_i - b \|)\frac{\vec{x}_i - b}{\| \vec{x}_i - b \|}, \qquad (16)$$

where $b$ is the closest point on the boundary to point $\vec{x}_i$, and the functional form is taken to be $f(d) = f_{\text{ster}}(1 - d/\ell_{\text{ster}})1_{d \le \ell_{\text{ster}}}$. That is, the steric force occurs when the point on the nucleolus surface is within a distance $\ell_{\text{ster}}$ of a corresponding boundary. The heterochromatin thickness (either surface or pillar) measured from experiments is used for $\ell_{\text{ster}}$.

For each pillar radius, heterochromatin is populated with a fixed thickness around all non-pillar surfaces, and then around pillar surfaces and the nucleolus itself with a radius-varying thickness.

In the main text, the generated geometries use force-magnitude parameters $f_{\text{ster}} = 1$, $f_{\text{area}} = 5$, $f_{\text{elast}} = 5$, chosen to qualitatively match shapes of nucleoli to those seen in experimental images. For reproducibility, other parameters taken were $W = 15$, $H = 10$ (to generate geometries), $H = 5$ (for diffusion model), and the center initial circle of the nucleolus is placed at $[0,4]$. The resulting geometries can be seen in ***Figure S8A.***

***Supplementary References***


1. Politz JCR, Tuft RA, Pederson T: Diffusion-based Transport of Nascent Ribosomes in the Nucleus. *Molecular Biology of the Cell* 2003, 14:4805-4812.
2. Ruland J, Kruger A, Dorner K, Bhatia R, Wirths S, Poetes D, Kutay U, Siebrasse J, Kubitscheck U: Nuclear export of the pre-60S ribosomal subunit through single nuclear pores observed in real time. *Nature Communications* 2021, 12.
3. Leech V, Hazel JW, Gatlin JC, Lindsay AE, Manhart A: Mathematical modeling accurately predicts the dynamics and scaling of nuclear growth in discrete cytoplasmic volumes. *Journal of Theoretical Biology* 2022, 533:110936.
4. Lindsay AE, Bernoff AJ, Ward MJ: First Passage Statistics for the Capture of a Brownian Particle by a Structured Spherical Target with Multiple Surface Traps. *Multiscale Modeling & Simulation* 2017, 15:74-109.
5. Lawley SD, Keener JP: A New Derivation of Robin Boundary Conditions through Homogenization of a Stochastically Switching Boundary. *SIAM Journal on Applied Dynamical Systems* 2015, 14:1845-1867.
6. Junod SL, Tingey M, Kelich JM, Goryaynov A, Herbine K, Yang W: Dynamics of nuclear export of pre-ribosomal subunits revealed by high-speed single-molecule microscopy in live cells. *iScience* 2023, 26:107445.
7. Landvogt L, Ruland JA, Montellese C, Siebrasse JP, Kutay U, Kubitscheck U: Observing and tracking single small ribosomal subunits in vivo. *Methods* 2019, 153:63-70.
8. Copos CA, Guy RD, Walcott S, Alamo JCd, Mogilner A: Mechanosensitive Adhesion Explains Stepping Motility in Amoeboid Cells. *Biophysical Journal* 2017, 112:433a.
